# Sugar signaling modulates SHOOT MERISTEMLESS expression and meristem function in Arabidopsis

**DOI:** 10.1101/2023.01.08.522175

**Authors:** Filipa L. Lopes, Pau Formosa-Jordan, Alice Malivert, Leonor Margalha, Ana Confraria, Regina Feil, John E. Lunn, Henrik Jönsson, Benoît Landrein, Elena Baena-González

## Abstract

In plants, development of all above-ground tissues is controlled by the shoot apical meristem (SAM) which balances cell proliferation and differentiation to allow life-long growth. To maximize fitness and survival, meristem activity is adjusted to the prevailing conditions through a poorly understood integration of developmental signals with environmental and nutritional information. Here, we show that sugar signals influence SAM function by altering the protein levels of SHOOT MERISTEMLESS (STM), a key regulator of meristem maintenance. STM is less abundant in the inflorescence meristems of plants grown or treated under limiting light conditions, with lower STM levels correlating with lower sugar content in these meristems. Additionally, sucrose but not light is sufficient to sustain STM accumulation in excised inflorescences. Plants overexpressing the α1-subunit of SUCROSE-NON-FERMENTING1-RELATED KINASE 1 (SnRK1) accumulate less STM protein under optimal light conditions, despite higher sugar accumulation in the meristem. Furthermore, SnRK1α1 interacts physically with STM, suggesting a direct local repression. Surprisingly, silencing *SnRK1α* in the meristem leads to reduced *STM* expression and severe developmental phenotypes previously associated with STM loss-of-function. Altogether, we demonstrate that sugars promote STM accumulation and that the SnRK1 sugar sensor plays a dual role in the SAM, limiting STM abundance under unfavorable conditions but being required for overall meristem organization and integrity. This highlights the importance of sugars and SnRK1 signaling for the proper coordination of meristem activities.

## INTRODUCTION

Unlike animals, plants generate organs throughout their life cycle. Postembryonic development relies on stem cell reservoirs localized in specialized tissues known as meristems. The shoot apical meristem (SAM) is the site where above-ground organogenesis is initiated, giving rise to leaves, axillary buds, flowers, and the stem. The SAM is organized into functionally distinct subdomains in which cell division and differentiation are tightly coordinated to maintain the integrity and regenerative potential of the meristem. The concerted regulation of the different regions requires the activity of several transcriptional networks and hormone signaling pathways (1).

In Arabidopsis (*Arabidopsis thaliana*), SAM homeostasis is finely controlled by a negative-feedback loop between WUSCHEL (WUS) and CLAVATA3 (CLV3) (2, 3). WUS is a mobile transcription factor produced in the organizing center that underlies the stem cell layers. WUS moves through plasmodesmata into the overlying stem cells to promote their proliferation and maintain pluripotency (4, 5). In addition, WUS induces stem cells to produce the CLV3 peptide, which in turn diffuses into the organizing center, where it inhibits *WUS* expression. The WUS-CLV3 feedback loop therefore enables a dynamic adjustment of the size of the stem cell pool (6).

Meristematic activity is also regulated by members of the THREE-AMINO-ACID-LOOP-EXTENSION (TALE) homeodomain protein superfamily which includes KNOTTED1-like homeobox (KNOX) and BEL-like homeobox (BLH or BELL) transcription factors. One key regulator of the TALE family is SHOOT MERISTEMLESS (STM), which is essential both for SAM establishment and SAM maintenance (7). Unlike WUS, STM is expressed throughout the SAM except at the sites of initiating primordia (7, 8). STM suppresses differentiation independently of WUS and promotes cell division by inducing the expression of *CYCLIN D3* (*CYCD3*) and *ISOPENTENYL TRANSFERASE7* (*IPT7*) (9), which encodes a key enzyme involved in cytokinin (CK) biosynthesis (10). CKs, in turn, are involved in stem cell maintenance, influencing SAM size and organ production through WUS and STM (11–13).

Loss-of-function *stm* mutants exhibit defects in SAM formation and maintenance, leading to growth arrest at the seedling stage due to exhaustion of stem cells (7). In addition, the most severely affected mutants like *stm-1* display fusions of cotyledons and other organs, indicating a role for STM also in boundary specification (7, 14, 15). Indeed, incipient organ primordia are formed at sites where *STM* expression is low, coincident with auxin accumulation and the expression of transcription factors (TFs) that promote organ formation and repress *STM*, including ASYMMETRIC LEAVES1 and 2 (AS1 and AS2) and members of the TEOSINTE BRANCHED1/CYCLOIDEA/PCF1 (TCP) family (16–20). In the inflorescence meristem, *STM* expression is also enhanced in boundaries, notably in response to mechanical forces, and is required for correct boundary folding (21).

Recent work revealed that STM heterodimerizes with WUS, enhancing WUS binding to the *CLV3* promoter and *CLV3* expression, and repressing stem cell differentiation. Conversely, WUS is required for the expression of *STM*, which thereby enhances WUS-mediated stem cell activity (22). STM is also regulated through an interaction with BELL1-like homeodomain (BLH) proteins (23) and the formation of these heterodimeric complexes is essential for STM nuclear localization and, thus, proper function (24). STM interacts with the BLH proteins PENNYWISE (PNY), POUNDFOOLISH (PNF), and ARABIDOPSIS THALIANA HOMEOBOX GENE1 (ATH1) (24–27) which contribute redundantly with STM to meristem initiation and maintenance (28–30).

Because of their sessile lifestyle, plants continuously adjust their development to changes in the environment, and this is reflected in the dynamic nature of the SAM. In addition to its maintenance by a network of TFs and hormonal signals, the SAM also responds to environmental signals that influence the relative size of its subdomains and the type and number of organs it produces. One of the external factors that affect meristem activity is light, which can exert a direct effect through photoreceptor-mediated signaling and an indirect effect by driving photosynthesis and sugar production (31–33). Both light and metabolic signals activate the TARGET OF RAPAMYCIN (TOR) protein kinase, which in turn promotes cell proliferation in the SAM *via* an increase in the expression of S-phase genes (34, 35). In addition, TOR induces *WUS* expression partially through an effect on CK degradation (33, 36).

TOR activity is often antagonized by SUCROSE NON-FERMENTING1-RELATED KINASE 1 (SnRK1) which, like TOR, translates environmental information into metabolic and developmental adaptations (37–39). SnRK1 is a heterotrimeric protein kinase complex, composed of an α-catalytic subunit and two regulatory β- and γ-subunits. In Arabidopsis, the α-subunit is present in two major isoforms, SnRK1α1 and SnRK1α2 (also known as KIN10 and KIN11). The SnRK1 complex is activated under low carbon conditions to promote energy saving and nutrient remobilization strategies, whilst TOR is activated in response to nutrient abundance to promote cell proliferation and growth adaptations (37–39).

Despite the importance of STM for establishing and maintaining SAM function and our increasing understanding of how STM activity is controlled by other transcriptional regulators, it is unknown if environmental signals modulate STM expression and/or activity. Here, we make use of plants expressing transcriptional and translational STM reporters to investigate this question. We show that STM protein accumulation does not respond to CK but is clearly induced by photosynthesis-derived sugars. We also show that the SnRK1 sugar sensing kinase is active in the SAM and that it is involved in adjusting STM protein levels to the light conditions. Finally, we show that SnRK1 is also required to maintain *STM* expression, meristem organization and integrity.

## RESULTS

### Light promotes STM protein accumulation

Light is essential for proper plant development and physiology. To investigate a potential regulatory role of light on STM levels, we made use of an Arabidopsis (Col-0) reporter line in which a fluorescently-tagged form of the STM protein (STM-VENUS) is expressed under the control of the *STM* promoter *[pSTM::STM-VENUS* (21, 40)]. We measured STM-VENUS levels in inflorescence meristems from 5-week-old plants grown under, or transiently treated with different light conditions. In one set of experiments, we compared STM-VENUS levels between plants grown under two different irradiances [60 *vs*. 170 μmol m^-2^ s^-1^, referred to as low light (LL) and high light (HL), respectively]. Irradiance had a strong impact on STM accumulation, with the mean STM-VENUS levels of plants grown under LL being 76% of those grown under HL (Fig. 1A-B). In a second set of experiments, we compared STM-VENUS levels between HL-grown plants transferred to darkness for up to 72 h and their corresponding controls maintained under HL conditions. Incubation under darkness had a very severe impact on STM accumulation, with STM-VENUS levels of plants subjected to a 72 h dark treatment being 39% of those prior to the treatment (Fig. 1A, 1C).

**Figure 1.**
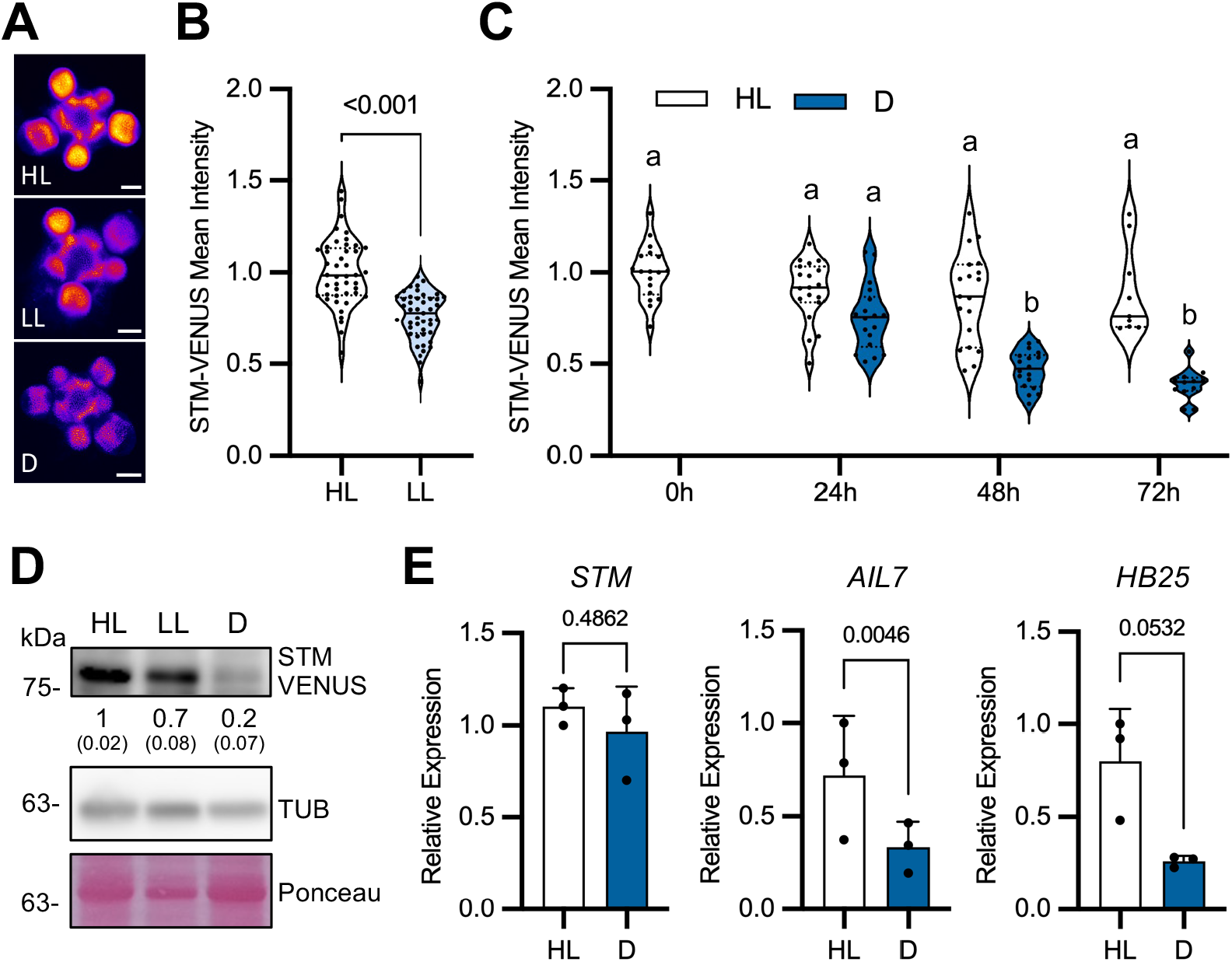
Effect of light on STM expression. **A-C,** STM-VENUS expression in SAMs of *pSTM::STM-VENUS* plants grown under high light (HL) or low light (LL) conditions or transferred from HL to darkness (D) or kept under HL for the indicated times. **A**, Representative STM-VENUS images of SAMs from HL and LL-grown plants and of plants transferred to D for 48 h. Scale bar, 50 μm. **B, C**. Quantification of STM-VENUS signal. **B,** Plots show SAM measurements of plants grown as 3 independent batches normalized by the mean of the HL condition of each batch (HL, *n*=44; LL, *n*=45). Student’s *t*-test (*p*-value shown). **C,** Plots show SAM measurements of plants grown as 2-3 independent batches normalized by the mean of the HL condition of each batch (0h, *n*=18; 24 h L, *n*=19; 24 h D, *n*=18; 48 h L, *n*=19; 48 h D, *n*=18; 72 h L, *n*=9; 72 h D, *n*=12). Different letters indicate statistically significant differences (Kruskal-Wallis with Dunn’s test; *p*<0.05). **D,** Immunoblot analyses of STM protein levels in SAMs of *pSTM::STM-VENUS* plants grown under HL or LL conditions or grown in HL and transferred to D for 48 h. TUBULIN (TUB) and Ponceau staining serve as loading controls. Numbers refer to mean STM-VENUS amounts in LL and D as compared to HL (*n*=2; each a pool of 5 SAMs; in parenthesis, SEM). **E,** RT-qPCR analyses of *STM* and STM target genes *AIL7* and *HB25* in SAMs of *pSTM::STM-VENUS* plants grown in HL and transferred to D or kept in HL for 48 h. Graphs show the average of 3 independent samples, each consisting of a pool of 5 SAMs. Paired ratio *t-* test (*p*-values shown).

To assess whether the impact of light on STM levels was a general effect on protein abundance in meristems, we extracted total proteins from SAMs of plants constantly grown under HL, LL, or treated with 48h of darkness and compared STM levels to those of the housekeeping protein TUBULIN (TUB) by immunoblotting (Fig. 1D). These analyses confirmed the microscopy results regarding STM-VENUS accumulation, showing that, in LL and dark-treated plants, STM levels were 71% and 23%, respectively, of the STM levels in HL. The immunoblots revealed no impact of the light conditions on TUB accumulation, indicating that the lower STM levels were not caused by a general decrease in protein accumulation. Finally, to assess if low STM accumulation could be due to reduced *STM* transcript abundance, we dissected SAMs of plants kept under HL conditions or subjected to 48 h darkness and analyzed *STM* transcript levels by quantitative RT-PCR (qPCR). *STM* levels were not significantly affected by the dark treatment (Fig. 1E), showing that the differences in protein accumulation are not due to changes in *STM* transcription or transcript stability. On the other hand, the levels of *AINTEGUMENTA-LIKE 7* (*AIL7*) and *HOMEOBOX PROTEIN 25* (*HB25*), known gene targets of STM (41), were reduced upon the dark treatment (Fig. 1E). This is also consistent with the lower STM-VENUS abundance and indicates decreased STM activity in the SAM in these conditions.

### The response of STM to light is CK-independent and involves sugars

Several lines of evidence suggest that, like *WUS, STM* expression could be directly regulated by CK (11, 42). To investigate whether CK could also regulate STM at the protein level and hence be involved in the response of STM to light, we first tested whether light influenced CK signaling in inflorescence meristems. To this end we used plants expressing the synthetic CK reporter *pTCSn::GFP* (43) in similar experiments as described for the *STM-VENUS* reporter line. In plants grown in LL or subjected to a 48 h dark treatment,*pTCSn::GFP* levels were 74% (Supplementary Fig. S1A-B) and 46% (Supplementary Fig. S1A, S1C) of those in HL plants, respectively. These observations show that CK signaling in inflorescence meristems is, like in vegetative meristems (33) affected by light. We next examined whether CK could impact STM levels in inflorescence meristems. For this, we excised SAMs of HL-grown *STM-VENUS* plants and maintained them under HL *in vitro* (13) for different periods of time in the absence or presence of 500 nM 6-benzylaminopurine (BAP), a synthetic CK. Dissection of the SAMs led to a strong reduction of the STM-VENUS (Supplementary Fig. S1D) and the *pTCSn::GFP* (Supplementary Fig. S1E) reporter signals, as previously described for the *pTCSn::GFP* and *pWUS::GFP* reporters (13). However, in contrast to *pTCSn::GFP* [Supplementary Fig. S1E; (13)], CK could not sustain STM-VENUS levels (Supplementary Fig. S1D), indicating that the effect of light on STM-VENUS is likely CK-independent.

Light plays direct signaling functions through various photoreceptors but also signals indirectly through sugars produced by photosynthesis. We therefore wondered whether the effect of light on STM was direct or mediated by sugars. To investigate this, we first measured the levels of sucrose, glucose, and fructose in the rosettes (Supplementary Fig. S2A) and SAMs (Fig. 2A) of HL- and dark-treated plants. We also measured the levels of Tre6P, a regulatory sugar-phosphate that serves as a signal of the plant sucrose status and that is crucial for sucrose homeostasis, growth promotion, and developmental progression (44). In the light, the levels of sucrose, glucose, and Tre6P were, respectively, 2.1-, 2.2-, and 7.9-fold higher in the SAM than in the rosette, whilst fructose accumulated to comparable levels in the two organs (Fig. 2A and Supplementary Fig. S2A). Incubation in the dark led to a marked depletion of sucrose and fructose both in rosettes (15% and 11% of the levels in the light, respectively) and SAMs (8% and 4% of the levels in the light, respectively), with a much milder reduction being observed for glucose, the most abundant sugar in the SAM (44% and 35% of the levels in the light in rosettes and SAMs, respectively). Tre6P levels were also much lower in dark-treated plants (11% and 3% of the levels in the light in rosettes and SAMs, respectively), reflecting the drop in sucrose levels. To further distinguish between a light and a sugar effect, we excised inflorescences at around 3 cm from the apex and placed them for 48 h in liquid medium. Similarly to what was observed in dissected SAMs (Supplementary Fig. S1D), STM-VENUS signal decreased markedly in cut inflorescences as compared to the uncut controls (Fig. 2B). Furthermore, light alone was not sufficient to sustain STM-VENUS expression, as the signal was comparable in cut inflorescences incubated in the light and in the dark (47% and 43% of the levels in the uncut control, respectively). These results, obtained with a double reporter line (*pSTM::STM-VENUS/pSTM::TFP-N7;* Fig. 2B), were similar to those obtained for plants expressing STM-VENUS alone (Supplementary Fig. S2B).

**Figure 2.**
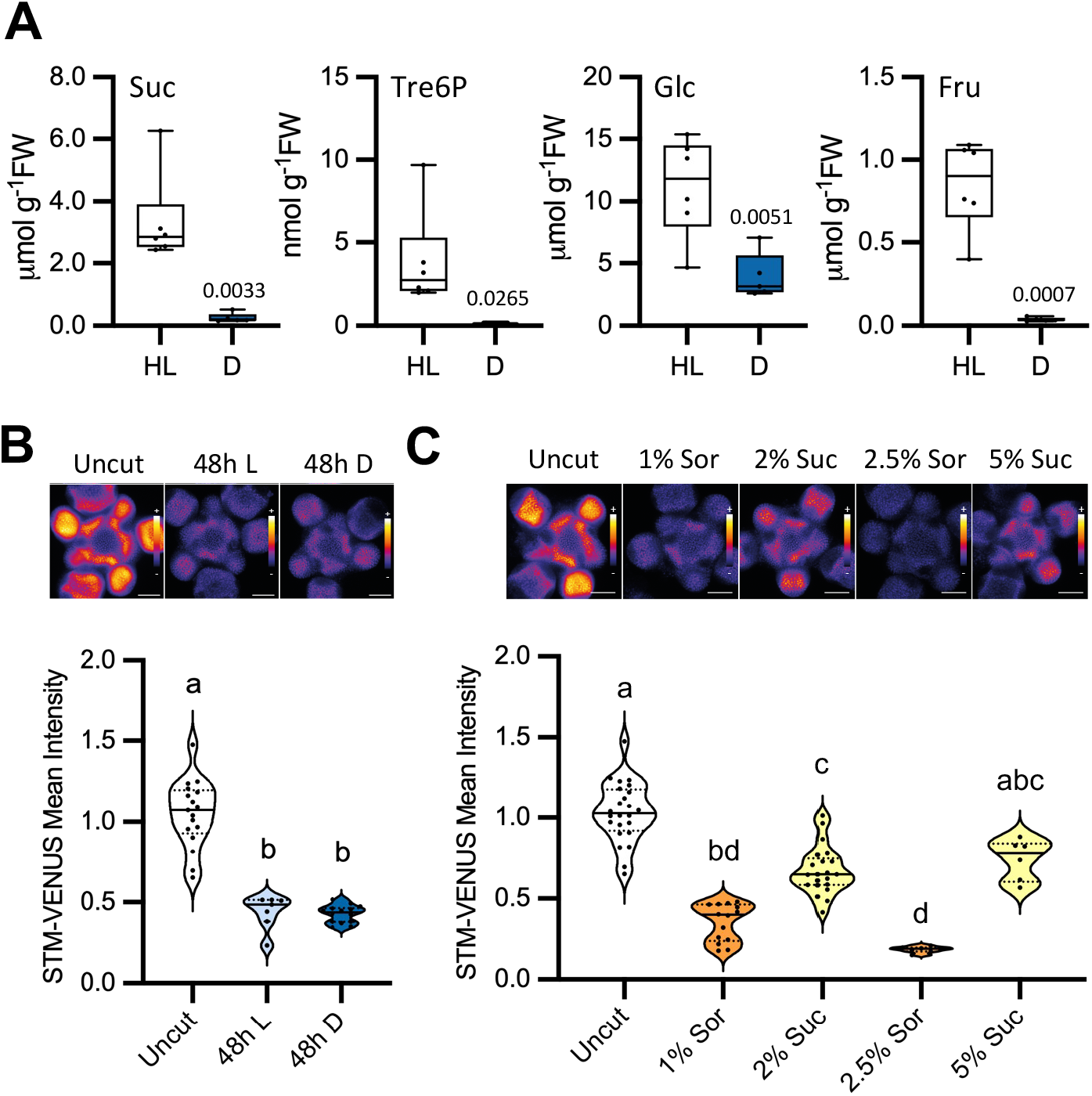
Effect of sugars on STM levels. **A,** Effect of light on the levels of soluble sugars in SAMs of *pSTM::STM-VENUS* plants grown in high light (HL) and transferred to darkness (D) or kept in HL for 48 h. Suc, sucrose; Tre6P, trehalose 6-phosphate; Glc, glucose; Fru, fructose. Plots show measurements of 5-6 samples, each consisting of a pool of 5 SAMs from plants grown as 2 independent batches. Welch’s *t*-test (*p*-value shown). **B,** Effect of light on STM-VENUS levels in cut inflorescences. Inflorescences of *pSTM::STM-VENUS/pSTM::TFP-N7* plants grown under HL were cut and placed in medium without sugar for 48 h under HL (L) or dark (D) conditions, after which the SAMs were dissected and imaged (VENUS). Upper panel, representative STM-VENUS images of SAMs. Scale bar, 50 μm. Lower panel, plots showing SAM measurements of plants grown as 1-2 independent batches normalized by the mean of the uncut condition of each batch (uncut, *n*=31, 2 batches; 48 h L, *n*=14, 1 batch; 48 h D, *n*=21, 2 batches). Different letters indicate statistically significant differences (Kruskal-Wallis with Dunn’s test; *p*<0.05). **C,** Effect of sugar on STM-VENUS levels in cut inflorescences. Inflorescences of *pSTM::STM-VENUS/pSTM::TFP-N7* plants grown under HL condition were cut and placed under darkness for 48 h in medium with sucrose (Suc; 2% and 5%) or sorbitol (Sor; 1% and 2.5%) as osmotic control. SAMs were thereafter dissected and imaged (VENUS). Upper panel, representative STM-VENUS images of SAMs. Scale bar, 50 μm. Lower panel, plots showing SAM measurements of plants grown as 1-3 independent batches normalized by the mean of the uncut condition of each batch (uncut, *n*=31, 3 batches; 1% Sor, *n*=21, 2 batches; 2% Suc, *n*=28, 3 batches; 2.5% Sor, *n*=7, 1 batch; 5% Suc, *n*=6, 1 batch). Different letters indicate statistically significant differences (Kruskal-Wallis with Dunn’s test; *p*<0.05). The same batches of HL-grown uncut plants served as controls for the experiments shown in (**B)** and (**C)**.

To test if the underlying cause was sugar deprivation, we first incubated excised inflorescences of the double marker line for 48 h in darkness in medium supplemented with increasing concentrations of sucrose. Sorbitol, which is not a readily metabolized carbon source, was used as an osmotic control. Sucrose was able to sustain STM-VENUS accumulation, and its effect was dose-dependent, leading to STM-VENUS levels close to those of uncut inflorescences when supplied at a 5% concentration (72% of the uncut control values as compared to 18% in the corresponding 2.5% sorbitol control; Fig. 2C). STM-VENUS levels did not increase in response to sorbitol, indicating that the effects of sucrose were not osmotic. Similar results were obtained for the single STM-VENUS reporter line (Fig. S2C). To test if the observed effects were transcriptional, we monitored the activity of the *pSTM::TFP-N7* transcriptional reporter. Quantification of the *pSTM::TFP-N7* signal revealed no significant repression of *STM* promoter activity upon inflorescence excision and incubation in darkness (Supplementary Fig. S2D), consistent with the results obtained by qRT-PCR in intact plants (Fig. 1E). In addition, incubation in sucrose or sorbitol-containing media had minor effects on TFP levels (Supplementary Fig. S2E) as compared to STM-VENUS (Fig. 2C), with the most severe condition (2.5% sorbitol) leading to 65% of the signal in the uncut control as compared to the 18% of the equivalent STM-VENUS samples. This indicates that the effect of sugar deprivation on STM levels does not rely on transcriptional regulation of *STM*.

### The SnRK1 sugar sensor is expressed in the SAM and influences STM levels

One major component of sugar signaling is the SnRK1 protein kinase, that is activated under conditions of low energy and is conversely repressed by sugars (45). Given its well-established role as a sugar sensor and the increasing number of studies implicating SnRK1 in developmental processes (37, 39), we next investigated whether SnRK1 could be involved in the regulation of SAM function through STM. To this end, we used a line expressing SnRK1α1-GFP under the control of the *SnRK1α1* promoter and other gene regulatory regions (46). A clear SnRK1α1-GFP signal was detected in the SAM, showing a stronger intensity in the peripheral regions, and developing organs (Fig. 3A). To further confirm this expression and to assess whether SnRK1α1 might be enriched in the SAM relative to other organs of the plant, we extracted total proteins from rosettes and shoot apices of 6 to 7-week-old plants and compared the relative levels of SnRK1α1 by immunoblotting (Fig. 3B). For the same amount of total protein, shoot apices contained higher amounts of SnRK1α1 suggesting that SnRK1 is relatively more abundant in the SAM than in rosette leaves.

**Figure 3.**
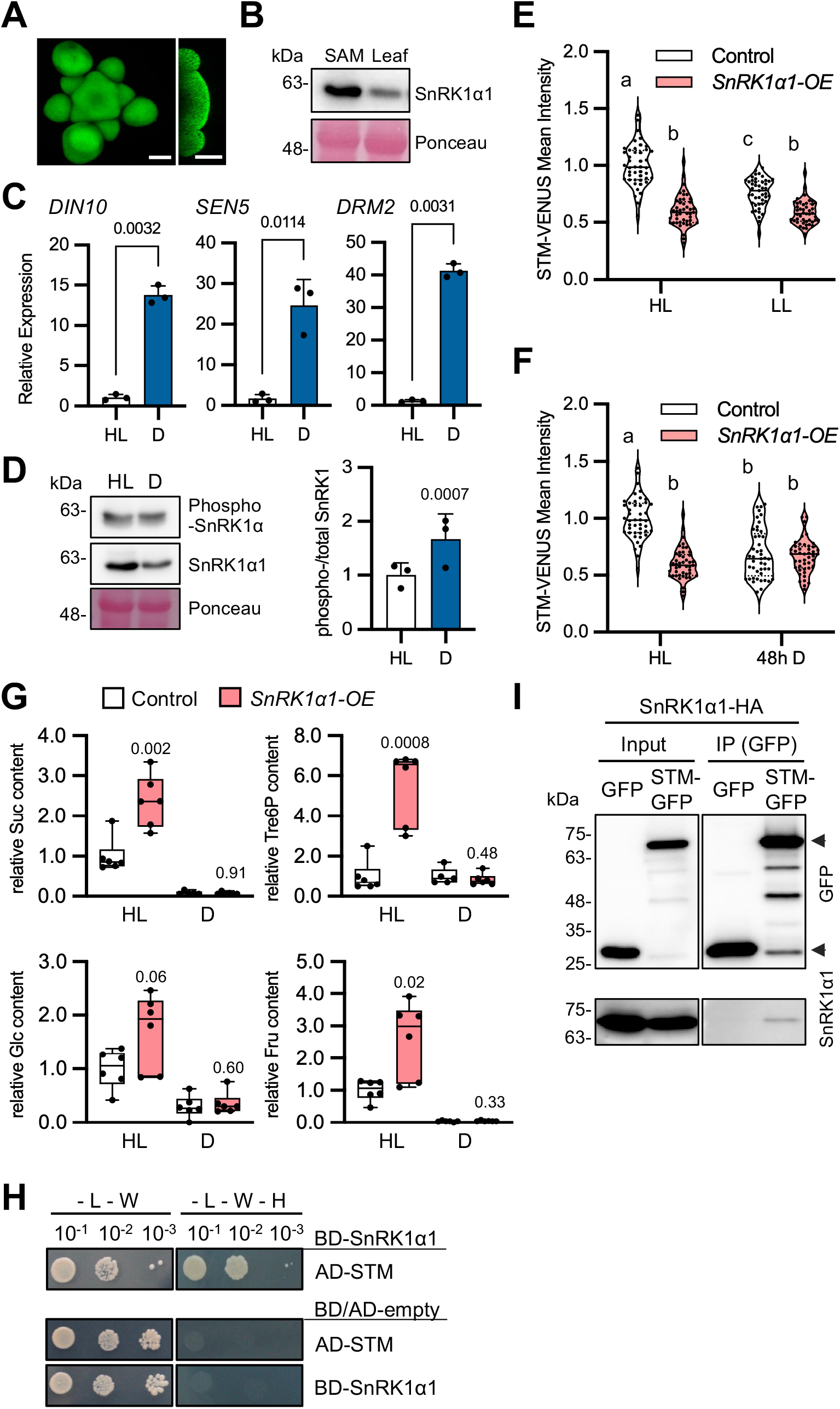
SnRK1 is expressed in the SAM and affects STM response to light. **A,** SnRK1α1-GFP imaging in the SAM. Right panel, SAM longitudinal section. Scale bars, 50 μm. **B,** Immunoblot analyses of SnRK1α1 in SAMs and rosette leaves of *pSTM::STM-VENUS* plants grown under high light (HL). 35 μg of total protein was loaded from SAM and leaf extracts. Ponceau staining serves as loading control. Similar results were obtained from two independent experiments. **C,** RT-qPCR analyses of SnRK1 signaling marker genes (*DIN10, SEN5, DRM2*) in SAMs of *pSTM::STM-VENUS* plants grown in HL and transferred to darkness (D) or kept in HL for 48 h. Graphs show the average of 3 independent samples, each consisting of a pool of 5 SAMs. Paired ratio *t-*test (*p*-values shown). **D,** Left, representative immunoblot of SnRK1α1 T-loop phosphorylation in SAMs of the plants described in **(C),** using antibodies recognizing the T175 phosphorylation (phospho-SnRK1α) or the total SnRK1α1 protein. Right, quantification of the mean SnRK1α phosphorylation (phospho-SnRK1α/total SnRK1α1) in D as compared to the ratio in HL (*n*=3; each a pool of 5 SAMs). Paired ratio *t-*test (*p*-value shown). **E-F,** STM-VENUS expression in SAMs of control and *SnRK1α1-OE* plants grown under HL or low light (LL) conditions (**E**) or grown under HL and transferred to darkness (D) or kept under HL for 48 h (**F**). The same batches of HL-grown plants served as controls for the experiments shown in (**E)** and (**F)**. HL and LL-grown STM-VENUS samples are replotted from Fig. 1B as a reference. Plots show SAM measurements of plants grown as 3 independent batches normalized by the mean of the HL condition of each batch (control, HL: *n*=44, LL: *n*=45, D: *n*=45; *SnRK1α1-OE*, HL: *n*=45, LL: *n*=45; D: *n*=45). Different letters indicate statistically significant differences (Kruskal-Wallis with Dunn’s test; *p*<0.05). **G,** Effect of light on the levels of sugars in SAMs of *SnRK1α1-OE* plants as compared to the control. Suc, sucrose; Tre6P, trehalose 6-phosphate; Glc, glucose; Fru, fructose. Plots show measurements of 5-6 samples, each consisting of a pool of 5 SAMs from plants grown as 2 independent batches. Welch’s *t*-test (mutant *vs*. control for each condition; *p*-values shown). **H**, Yeast-two hybrid assays examining the interaction of SnRK1α1 with STM. Protein interaction was determined by monitoring yeast growth in medium lacking Leu, Trp and His (-L-W-H) compared with control medium only lacking Leu and Trp (-L-W). Upper panel, yeast growth in cells co-expressing AD-STM, with BD-SnRK1α1. Lower panel, negative controls of yeast transfected with the indicated AD/BD-constructs and the complementary BD/AD-empty vectors. BD and AD, DNA binding and activation domains of the GAL4 transcription factor, respectively. Increasing dilutions of transformed yeast cells are shown (10^-1^, 10^-2^, 10^-3^). Experiments were performed three times with similar results. **I**, Co-immunoprecipitation (co-IP) experiments using Arabidopsis Col-0 mesophyll cell protoplasts co-expressing SnRK1α1-HA with STM-GFP or GFP alone. GFP-tagged proteins were immunoprecipitated and co-immunoprecipitation of SnRK1α1 was assessed by immunoblotting with an HA antibody. Arrowheads, STM-GFP (upper) and GFP (lower). Experiments were performed three times with similar results.

To test if SnRK1 is functional in the meristem, we used SAMs dissected from HL- or dark-treated plants (48 h) to measure the activity of the SnRK1 signaling pathway using the expression of downstream target genes (45) as a readout (Fig. 3C). Under control conditions, SnRK1-regulated genes were barely expressed, consistent with the pathway being inactive. However, after 48 h of darkness, a marked upregulation of these genes was observed (Fig. 3C), indicating an activation of SnRK1 signaling in the SAM. The induction of SnRK1-regulated genes in darkness was accompanied by a reduction in total SnRK1α1 levels (Fig. 3D), consistent with a tight coupling between SnRK1 activity and degradation (46), and by an increase in the relative phosphorylation of the SnRK1α1 (T-loop) that is essential for SnRK1 activity (45).

To investigate whether SnRK1 is involved in STM regulation, we introgressed the *pSTM::STM-VENUS* reporter construct into a line overexpressing SnRK1α1 *[35S::SnRK1α1*, hereafter referred to as *SnRK1α1-OE;* (47)] and monitored STM-VENUS levels in different light conditions. When plants were grown under HL, the levels of STM-VENUS in the *SnRK1α1-OE* were 60% of those in control plants (Fig. 3E), a decrease that could not be explained by differences in *STM* transcript levels (Supplementary Fig. S3). However, the differences between the two genotypes became smaller when plants were grown in LL (STM-VENUS levels in *SnRK1α1-OE* were 77% of those in HL plants; Fig. 3E) and negligible when subjected to a 48 h dark treatment (STM-VENUS levels in *SnRK1α1-OE* were 97% of those in control plants; Fig. 3F). STM-VENUS levels thus appeared to be constitutively low and largely insensitive to the light conditions in *SnRK1α1-OE* plants. This contrasted with control plants which, in response to restrictive light conditions, reduced STM-VENUS accumulation to levels equivalent to those of the *SnRK1α1-OE*. Lower STM-VENUS levels in the *SnRK1α1-OE* in HL could not be explained by lower sugar accumulation, as these plants had a higher content of sucrose, glucose, and fructose both in the SAM (Fig. 3G) and rosettes (Supplementary Fig. S4), although the differences were not always statistically significant due to the large variation of the *SnRK1α1-OE* samples. The levels of Tre6P, known to inhibit SnRK1 activity (48–50), were also markedly higher in the *SnRK1α1-OE* SAMs (5.6-fold) and rosettes (5-fold), consistent with previous observations in *SnRK1α1-OE* rosettes (51). During the dark treatment, however, all sugars were largely depleted, reaching similarly low levels in control and mutant samples.

Altogether these results suggest that SnRK1 is active in the SAM and that it contributes to the adjustment of STM protein levels, inhibiting STM accumulation when sugar levels decline. To further investigate the involvement of SnRK1 on STM regulation, we first used yeast-two-hybrid (Y2H) assays to test if SnRK1α1 can interact directly with STM (Fig. 3H). We observed that yeast co-expressing SnRK1α1 with STM were able to grow in selective medium but this was not the case when SnRK1α1 or STM were expressed individually with the corresponding empty vector controls, suggesting that these two proteins can interact. To determine if the SnRK1α1-STM interaction can also occur *in planta*, we next performed co-immunoprecipitation (co-IP) experiments using Col-0 mesophyll cell protoplasts expressing SnRK1α1-HA with STM-GFP or with GFP alone as a negative control. Immunoprecipitation with an anti-GFP antibody and subsequent Western blot analyses revealed that SnRK1α1 interacts with STM-GFP (Fig. 3I), but not with GFP alone, indicating that the interactions revealed by Y2H may also occur *in planta*. Taken together, our results suggest that the impact of SnRK1α1 on STM accumulation may be due to direct action of the SnRK1 kinase on STM in the meristem.

### Silencing *SnRK1α* in the SAM reduces STM expression and disrupts meristem function

To investigate further the possibility that SnRK1 acts locally in the meristem, we designed artificial microRNAs (amiRNAs) targeting both *SnRK1α1* and *SnRK1α2* in two different regions of the transcripts (*amiRα-1* and *amiRα-2*) and expressed these *amiRNAs* under the 5.7 kb promoter of *STM* in *STM-VENUS* plants (Fig. 4A). Immunoblot analyses confirmed a decrease in the activated form (phosphorylated in the T-loop) of SnRK1α in all lines, but this was accompanied by a decrease in total SnRK1α1 levels only in lines expressing *amiRα-2* (Fig. 4B).

**Figure 4.**
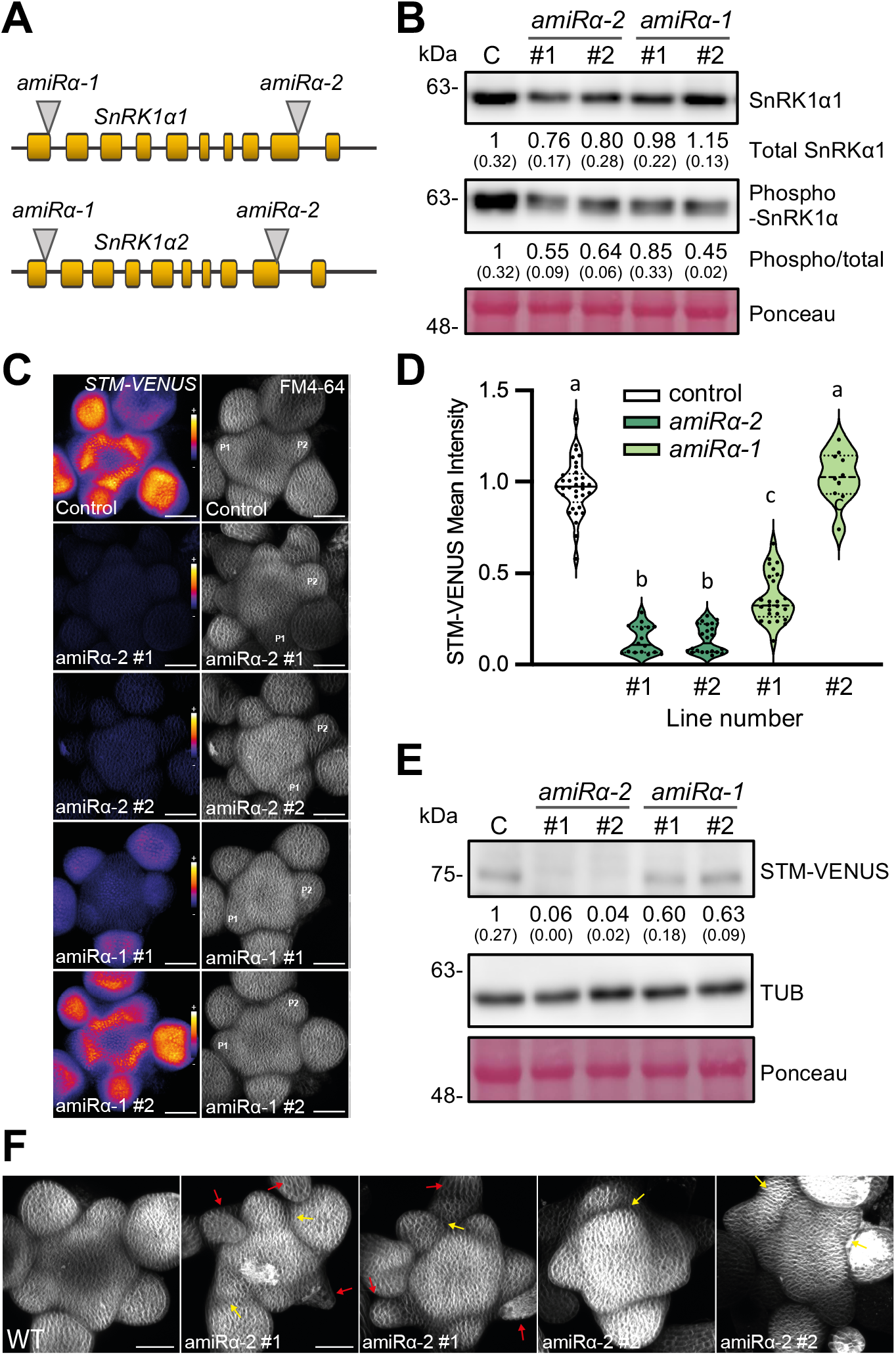
Silencing *SnRK1α* in the SAM compromises STM expression. **A.** Schematic localization of *amiRα-1* and *amiRα-2* target sites (grey triangles) in the *SnRK1α1* and *SnRK1α2* transcripts. Yellow blocks correspond to exons. **B.** Immunoblot analyses of SnRK1α T-loop phosphorylation in SAMs of plants expressing *pSTM::amiRα-1* (*amiRα-1*) or *pSTM::amiRα-2* (*amiRα-2*), using antibodies recognizing total SnRK1α1 or SnRK1α phosphorylated on T175 (phospho-SnRK1α). Ponceau staining serves as loading control. Numbers refer to mean SnRK1α1 amounts or mean SnRK1α phosphorylation (phospho-SnRK1α/total SnRK1α1) in the *amiRα* lines relative to the control *STM-VENUS* line (*n*=2; each a pool of 5 SAMs; in parenthesis, SEM). **C.** Representative meristems expressing STM-VENUS together with *pSTM::amiRα-1, pSTM::amiRα-2* or *pSTM::TFP-N7* as a control, and whose membranes were labelled with FM4-64. Left panels show the sum-slice projections of the STM-VENUS signal (color-coded using the Fire representation in ImageJ), and the right panels, the sum-slice projection of the FM4-64 signal. Scale bars, 50 μm*. P1* and *P2*, youngest visible and older flower primordia, respectively. **D**. Quantification of the STM-VENUS signal in the SAMs shown in (**C**). Plots show SAM measurements of plants grown as 2 independent batches (except *amiRα-1#2*, which grown as a single batch) normalized by the mean of the control line of each batch (control, *n*=34; *amiRα-2#1, n*=16; *amiRα-2#2, n=22; amiRα-1#1, n=25; amiRα-1#2 n*=10). Different letters indicate statistically significant differences (Kruskal-Wallis with Dunn’s test; *p*<0.05). **E.** Immunoblot analyses of STM-VENUS protein levels in SAMs of the *amiRα* lines as compared to the *STM-VENUS* control using antibodies recognizing STM. TUBULIN (TUB) and Ponceau staining serve as loading controls. Numbers refer to mean STM-VENUS amounts in the *amiRα* lines as compared to the control (*n*=2; each a pool of 5 SAMs; in parenthesis, SEM). **F.** Sum-slice projection of control line and *amiRα-2* showing additional defects in meristem organization. Red arrows point at bract-like structures while yellow arrows point at fusions between adjacent floral primordia. Scale bars: 50 μm.

To our surprise, depletion of SnRK1α resulted in decreased STM-VENUS accumulation, both at the protein (Fig. 4C-E) and transcript levels (Supplementary Fig. S5). The decrease in STM-VENUS levels was strongest in the lines with more compromised SnRK1α activity (*amiRα-2* lines) and weaker in the lines where the impact on SnRK1α activity was moderate or negligible (*amiRα-1* lines; Fig. 4B-E). Lower STM accumulation in the SAM of the *SnRK1α* amiRNA lines correlated with defects in SAM development, including altered phyllotaxy, reduced bulging and the appearance of bract-like structures in some floral meristems, as well as fusions between adjacent floral meristems (Fig. 4F). Organ fusions were also visible later in development and affected cauline and rosette leaves, petals, siliques, and stems (Fig. 5D,F). Consistent with a previous report (21), STM depletion caused defects in boundary formation, manifested as a decreased curvature at the boundary between the meristem and the incipient organ in the strong *amiRα-2* lines (Supplementary Fig. S6).

**Figure 5.**
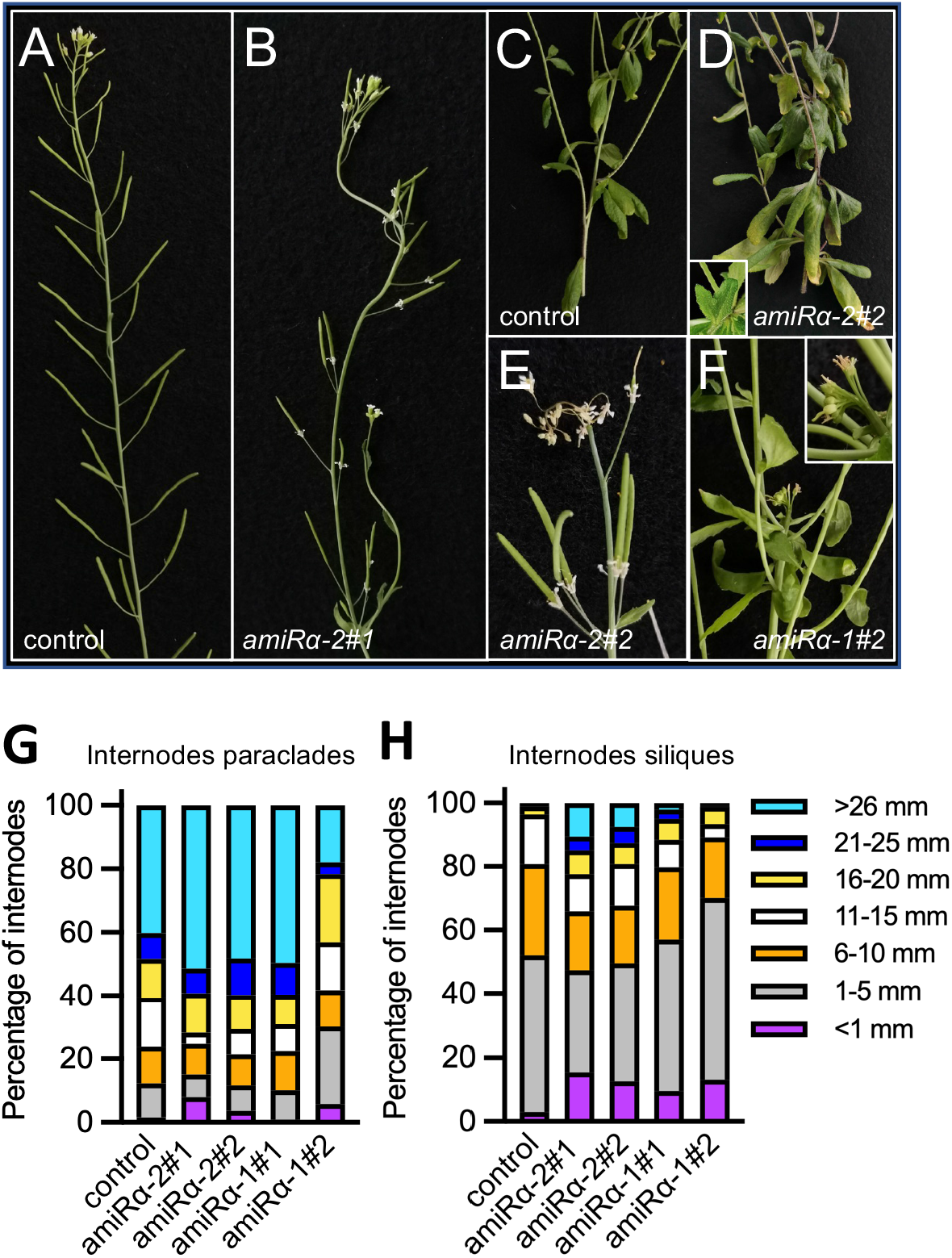
Silencing of *SnRK1α* in the SAM affects meristem function and plant architecture. **A-F,** Representative images of control (*STM-VENUS*, **A,C**) and *amiRα* (**B,D,E,F**) plants showing irregular internode length (**A,B**), clusters of leaves (**C,D,F**) and siliques (**A,B,E**), and termination of the main inflorescence (**F**) in the *amiRα* lines. Insets show organ fusion between leaves of an aerial rosette (**D**) and between pedicels and the stem (**F**). **G-H**, Quantification of the internode length defects in control and two independent lines of *amiRα-1* and *amiRα-2* mutants. Internode length was determined by measuring the length of the internodes between paraclades (**G**) and between the first 12 siliques (**H**), all counted acropetally. Internode length was scored in the indicated size ranges from the main inflorescence of 18 plants of each genotype. Graphs show the relative frequencies of each size class in the total number of internodes scored. All phenotypes were scored from plants grown under equinoctial conditions until the completion of flowering.

All *amiRα* lines displayed defects in internode elongation, with an increasing frequency of aberrantly long and aberrantly short internodes (Fig. 5A-B, G-H). Defects in internode elongation resulted in clusters of siliques (Fig. 5A,B,E) and what appeared to be aerial rosettes on the main inflorescence (Fig. 5C,D,F; Supplementary Fig. S7A; Table 1). These phenotypes have been linked to reduced *STM* expression and function (14, 52–54) and, accordingly, they were more severe in the *amiRα-2* plants, where the depletion of STM-VENUS is more pronounced (Fig. 4C-E). Plants expressing *amiRα-2* also exhibited reduced apical dominance with one or two axillary meristems often becoming activated well before flowering (39% and 28% of the *amiRα-2#1*, and *amiRα-2#2* plants, respectively; *n*=18; Supplementary Fig. S7B-C). A less frequent termination of the main meristem was also observed, after which, growth resumed from an axillary meristem (17% of *amiRα-2#1* plants; *n*=18; Supplementary Fig. S7C). Plants expressing *pSTM::amiRα* were also compared to the double reporter line as control (*pSTM::STM-VENUS/pSTM::TFP-N7*) to ensure that the observed phenotypes were not caused by the introgression of an additional *STM* promoter in the genome of the STM-VENUS line (Fig. 4C,D,F; Supplementary Fig. S7D).

**Table 1.**
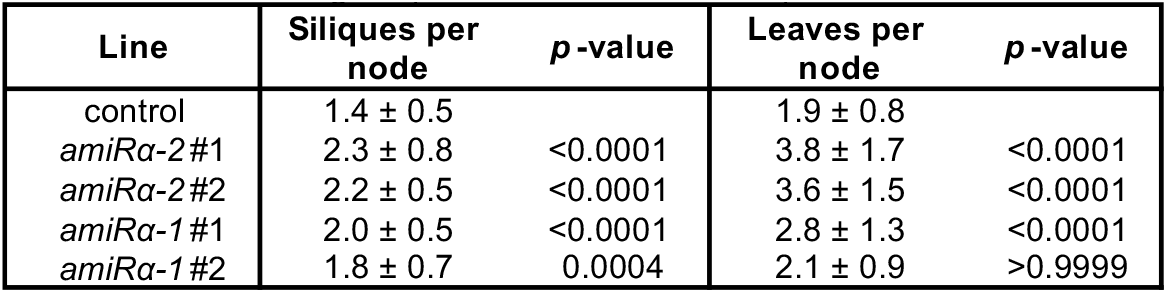
Number of organs per stem node in *amiRα* plants. Quantitative analyses of *amiRα* phenotypes. Measurements were taken from the main inflorescence of 18 plants of the *amiRα* lines and the *STM-VENUS* control line after flowering was completed. Numbers are averages and SD. *p*-values refer to differences between each mutant and the control (Kruskal-Wallis with Dunn’s test).

Collectively, these results indicate that SnRK1 plays critical functions in meristem organization and function and that this involves changes in *STM* expression.

## DISCUSSION

The capacity to generate organs throughout development is crucial for plant adaptation to the environment, enabling, amongst others, the replacement of leaves lost due to herbivory, the timing of growth to a specific season, or the switch to flower production when conditions are propitious for reproduction. However, how the SAM perceives environmental information and how this is translated into changes in meristem activity are poorly understood.

Here we show that light promotes STM accumulation through sugars. First, a clear correlation between STM-VENUS and SAM sugar levels was observed across different light conditions. STM-VENUS levels were lower in LL-grown or dark-treated plants than in plants grown and maintained under HL (Fig. 1A-C). A similar pattern was observed for sugar accumulation in the inflorescence meristems (Fig. 2A), consistent with a previous report on the impact of limiting photosynthetic rates, and thereby sugar supply to the sinks, on the growth and development of reproductive organs and meristem function (55). Second, STM-VENUS levels declined rapidly when inflorescences were excised from rosettes and this decline was similar in inflorescences maintained in the light or transferred to dark, showing that light is not sufficient to sustain STM levels in this system (Fig. 2B). The reason for this could be that light is sensed in leaves from which a light-related signal travels to the apex and that this remote light sensing and systemic communication is disrupted upon excision of the inflorescence. An alternative explanation is that the signal regulating STM levels is not light itself, but rather photosynthesis-derived sugars. The fact that the decline in STM-VENUS levels triggered by inflorescence excision could be largely suppressed by supplementing sucrose in darkness, argues that sucrose is sufficient to sustain STM-VENUS levels and that the effect of light observed in intact plants is indirect *via* photosynthesis and sugar production. The impact of sucrose on STM is in line with the reported effects of nutrients on *WUS* expression and on meristem function. Sugars contribute to meristem activation by inducing *WUS* in young seedlings (33) and nitrogen promotes *WUS* expression and meristem growth in the inflorescence *via* systemic CK signaling (13). However, in contrast to *WUS*, for which transcriptional regulation plays a major role (13, 33), we could not detect significant changes in *STM* transcript levels under our different growth conditions or treatments (Fig. 1E and Supplementary Fig. S3), supporting that STM is regulated at the protein level. Despite reports linking CK signaling to *STM* expression (11, 42), STM-VENUS levels did not increase in excised meristems treated with CK (Supplementary Fig. S1D). Even though we did not measure *STM* transcript accumulation under these conditions, this could mean that CK signals may influence STM more indirectly, *e.g*. by affecting *WUS* expression and stem cell number (56).

The rescue of STM-VENUS levels by sucrose in excised inflorescences suggests that sucrose is sensed locally in the meristem. This is in accordance with the enrichment and activity of the SnRK1 sugar sensor in the SAM (Fig. 3A-D) and with the importance of meristematic SnRK1 for SAM function (Fig. 4 and 5, Supplementary Fig. S6-7). Ubiquitous *SnRK1α1* overexpression caused a reduction in STM-VENUS levels under HL conditions (Fig. 3E) despite the high accumulation of soluble sugars and Tre6P in the rosettes and SAMs of the *SnRK1α-OE* plants (Fig. 3G). In addition, SnRK1α1 interacts physically with STM in yeast cells (Fig. 3H) and mesophyll cell protoplasts (Fig. 3I), supporting a close functional connection with STM and a local function for SnRK1 in the meristem.

Expressing *amiRs* targeting *SnRK1α1* and *SnRK1α2* under the control of the *STM* promoter demonstrated that SnRK1 acts locally in the SAM and is crucial for the proper coordination of meristem activities, with reduced SnRK1 activity causing a wide range of developmental defects (Fig. 5). The severity of the phenotypes caused by SnRK1α depletion contrasts with the apparent lack of developmental defects of the *SnRK1α-OE* line in our growth conditions. This is consistent with previous studies in which SnRK1α1 overexpression causes mostly developmental delays rather than defective organ development and arrangement (37, 39).

The finding that sucrose promotes STM-VENUS expression together with the fact that sugars repress SnRK1 activity may at first sight appear to conflict with the decline in STM-VENUS expression and abnormal meristem function caused by *SnRK1α* silencing. However, despite being generally considered a growth repressor, SnRK1 is also required for cell cycle progression (57) and for normal growth and development (58). Indeed, transient *SnRK1α1/α2* downregulation *via* virus-induced gene silencing leads to full growth arrest of plants (45) and double *snrk1α1 snrk1α2* null mutants could thus far not be recovered, suggesting that complete loss of SnRK1α is embryo lethal (59). A similar duality is observed for the AMP-activated protein kinase (AMPK), the homologue of SnRK1 in animals. Despite serving as a brake for cell proliferation through downregulation of TOR activity (60), AMPK is also essential for normal growth and development. For example, complete loss of the AMPKß1 subunit leads to cell cycle defects in neural stem and progenitor cells, causing profound abnormalities in brain development in mice (61). Along the same lines, hematopoietic stem cell function in mammals is disrupted both upon inactivation and overactivation of TOR signaling, indicating that a fine balance of this central regulator is required for coordinating proliferation, differentiation, and regeneration (62).

The effects of light, sucrose and SnRK1α1 overexpression on STM indicate that the underlying mechanisms do not rely on changes in *STM* transcript abundance (Fig. 1–3). Furthermore, the fact that the SnRK1α1 and STM proteins interact (Fig. 3H-I) suggests that the impact of SnRK1 on STM may be direct. The consequences of *SnRK1α* silencing, on the other hand, reveal a more complex scenario, involving reduced accumulation of both the STM transcript and protein, and severe developmental abnormalities. The phenotypes of the *amiRα* lines are highly reminiscent of those described for partial STM loss-of-function (14, 15, 21, 63), including organ fusions, altered phyllotaxy, defective internode elongation with clusters of siliques and leaves and, much less frequently, premature SAM termination (Table 1; Fig. 5; Supplementary Fig. S7). However, whether the impact of SnRK1α on STM in this case is direct or indirect through other factors requires further investigation. It is plausible that *SnRK1α* silencing causes hormonal imbalance and/or alterations in the cell cycle that feed back to *STM* expression.

Altogether, our work demonstrates that sucrose promotes STM accumulation and that this is counteracted by the SnRK1 sugar sensor, likely to adjust SAM activity to the environment (Fig. 6). Nevertheless, SnRK1 is also essential for meristem integrity, adding to the evidence that SnRK1 performs a dual function in the regulation of growth and that its activity needs to be finely balanced.

**Figure 6.**
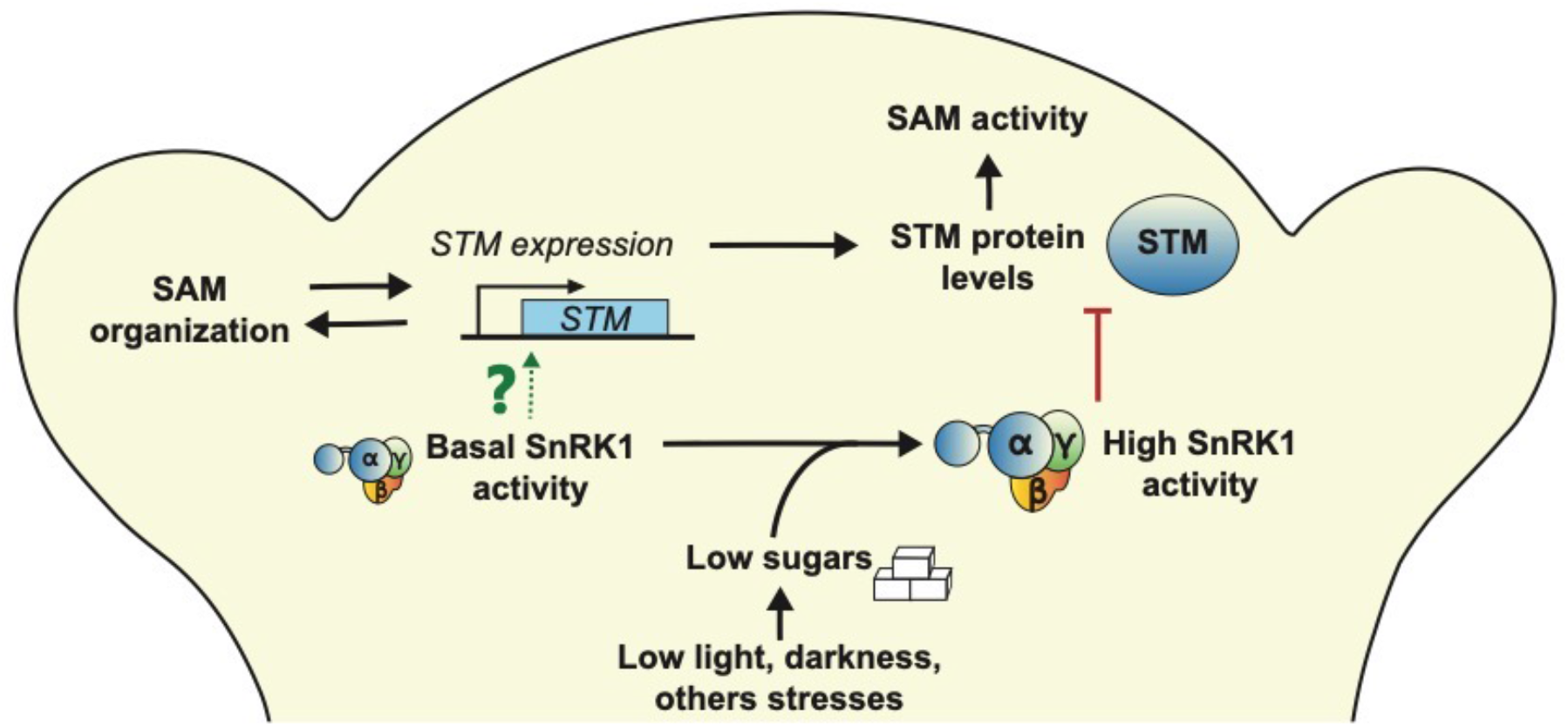
Model for the role of sugars and SnRK1 signaling in the SAM. Under favorable conditions, basal SnRK1 activity is required for meristem organization, with local *SnRK1α* silencing causing reduced *STM* expression and severe phenotypes related to SAM dysfunction. The mechanisms underlying these SnRK1 effects remain unknown (question mark). Under limiting light conditions or other unfavorable situations, sugar levels decrease, leading to a strong activation of SnRK1 signaling. This results in decreased STM protein accumulation, potentially through direct action of SnRK1α1 on STM to reduce SAM activity and growth.

## MATERIALS AND METHODS

A list of all primers, antibodies and plant lines used in this study is provided in Supplementary Table 1. Protein extraction and quantification, immunoblotting, RNA extraction, cDNA synthesis, qRT-PCR, sugar measurements, yeast-two-hybrid assays, and protoplast assays were carried out according to protocols described in *SI Materials and Methods*.

### Plant material

All *Arabidopsis thaliana* (L.) Heynh. plants used here are in the Columbia (Col-0) background. The *pSTM::STM-VENUS* line (*STM-VENUS*) was generated by transforming Col-0 plants with the plasmid described by Heisler and colleagues (13, 18). The *pTCSn::GFP* line was provided by Bruno Müller (43). The *SnRK1α1-GFP* [*pSnRK1α1::SnRK1α1-GFP::terSnRK1α1/snrk1α1-3*; (46)] and *SnRK1α1-OE [35S::SnRK1α1;* (47)] lines were previously described. For expression of STM-VENUS in the *SnRK1α1-OE* background, the *SnRK1α1-OE* and *STM-VENUS* lines were crossed, and homozygous progeny was selected on kanamycin and BASTA. For generating the *pSTM::STM-VENUS/pSTM::TFP-N7* line and the *pSTM::amiRα* lines, *STM-VENUS* plants were transformed with a construct to express TFP-N7 or an amiRNA targeting both *SnRK1α1* and *SnRK1α2 (amiRα-1* or *amiRα-2*) under the *STM* promoter (5.7 kb). Detailed descriptions of the cloning strategy and progeny selection are provided in *SI Materials and Methods*.

### Plant growth conditions

For most experiments, seedlings were initially grown in short-day conditions (8 h/16 h light/dark period) for 3-4 weeks and then transferred to continuous light (24 h light, temperature: 22°C) and kept under an irradiance of 60 (LL) or 170 (HL) μmol m^-2^ s ^-1^, provided by white fluorescent lamps. Unless otherwise indicated, plants were grown in HL conditions. For dark treatment, plants grown under HL were put into the dark for 24, 48 or 72 h. For the assays with excised inflorescences and STM-VENUS imaging of *amiRα* lines, seedlings were grown in short-day conditions (8 h/16 h light/dark period) for 3-4 weeks and then transferred to long-day conditions (16 h/8 h light/dark period; 22°C/18°C).

For experiments involving SAM imaging, gene expression or protein analyses, plants were grown in the indicated conditions until bolting, after which SAMs were dissected at the beginning of the flowering stage when the main inflorescence reached 3-5 cm in height.

For phenotyping the *amiRα* lines, seeds were germinated, and plants grown under equinoctial conditions (12 h/12 h light/dark period; 100-110 μmol m^-2^ s^-1^; 22°C/18°C). Phenotypes were scored when flowering was completed [stage 6.90; (64)].

### SAM imaging and quantification

For meristem imaging, the main inflorescence meristem of plants at the beginning of the flowering stage was cut 1-2 cm from the tip, dissected under a binocular stereoscopic microscope to remove all the flowers down to stage 3 [as defined in (65)] and transferred to a box containing Arabidopsis apex culture medium without sucrose (ACM: 2.2 g/l Duchefa Biochemie (www.duchefa-biochemie.com)-MS basal salt mixture with vitamins, pH adjusted to 5.8 with KOH, and 1.6% (w/v) agarose added).

For the time-lapse experiments examining the effect of exogenous CK, meristems were dissected from HL-grown plants and placed in a box of ACM with 1% (w/v) sucrose and 500 nM BAP or the equivalent volume of the BAP solvent (DMSO) as control. Meristems were thereafter returned to the constant HL cabinet for the indicated times and covered with water for imaging.

For the experiments examining the effect of exogenous sugar, inflorescences were dissected at about 3 cm from the apex from HL-grown plants at the beginning of the flowering stage and placed in a 2 mL Eppendorf tube containing 2 mL of liquid ACM (no sucrose) covered with parafilm pierced with a needle so that the inflorescence could be held in air while the base of the stem was submerged in the solution, supplemented with the indicated concentrations of sucrose or sorbitol as osmotic control. Sorbitol and sucrose were used at roughly equivalent concentrations, with 1% sorbitol and 2% sucrose corresponding to 54 mM and 58 mM, respectively. Inflorescences were thereafter kept in darkness inside the growth cabinet for the indicated times. Meristems were then dissected from the excised inflorescences, transferred to a box containing the same sucrose- or sorbitol-supplemented solid (1.6% agarose) medium and covered with water for imaging.

Meristems were imaged in water using a 20X long-distance water-dipping objective mounted either on a LSM880 (Zeiss; www.zeiss.com) or a SP8 (Leica; www.zeiss.com) confocal microscope. Z-stacks of 1-2 μm spacing were taken and the spacing was kept constant within a single experiment.

Confocal Z stacks were analyzed using the ImageJ software (https://fiji.sc/) and a custom-made code written in Matlab (Mathworks Inc., Natick, MA). ImageJ was used for generating the panel figures showing the fluorescence reporters in the SAMs. To generate the panels, z-projections (sum slices) of meristems expressing TFP, GFP or VENUS were performed, and the Fire color code was used to represent the signal. The expression levels of the different fluorescence reporters were analyzed by using the Matlab code (see https://gitlab.com/slcu/teamHJ/pau/RegionsAnalysis). *pTCSn::GFP* reporter was analyzed by using a previously described pipeline (13), which measures total fluorescence intensity of circular expression domains. STM-VENUS signal and TFP signal were analyzed by using a new pipeline that consists on the following. Firstly, a z-sum intensity projection is performed followed by a gaussian blur with a smoothing kernel with standard deviation sigma = 5 μm. Second, a region of interest (SAM core excluding the emerging floral organs) is drawn on the image projection and the mean intensity of the region is extracted. The mean intensity was chosen as a measure of the STM-VENUS and TFP levels instead of the total intensity to minimize the influence of manually drawing the region of interest and the new buds on the signal.

To measure the correlation between the folding of the boundary and the size of the primordia in *pSTM::amiRα-1* and *amiRα-2* lines, maximal projections were performed to outline the stage-2 primordia (65) and measure their projected area. Longitudinal sections passing through the middle of the primordia were also performed to measure the folding of the boundary using the angle tool of ImageJ as previously described (21). The relationship between primordia size and the folding of the boundary appeared to be more linear for smaller primordia. Therefore, only stage-2 primordia up to 4000 μm^2^ in size were considered for these measurements.

## ACKNOWLEDGEMENTS

We thank Vera Nunes for excellent plant management, Markus Heisler and Teva Vernoux for the *pSTM::STM-VENUS* constructs, and Bruno Müller for the *pTCSn::GFP* line. This work was supported by Fundação para a Ciência e a Tecnologia grants UIDB/04551/2020, UIDP/04551/2020, (GREEN-IT-Bioresources for Sustainability), PTDC/BIA-FBT/4942/2020, LISBOA-01-0145-FEDER-028128, PTDC/BIA-BID/32347/2017, PD/BD/128397/2017 (FLL) and by the Max Planck Society (RF, JEL, and PFJ), and the Gatsby Charitable Foundation (Grant GAT3395-PR4B to HJ). For the work carried in Lyon, we thank A. Lacroix, P. Bolland and J. Berger for technical assistance regarding plant cultivation; I. Desbouchages and H. Leyral for technical assistance regarding molecular biology work; C. Vial, L. Grangier, N. Camilleri and S. Maurin for administrative assistance; and the PLATIM facility of the SFR128 Biosciences for technical assistance regarding the microscopy.

## SUPPLEMENTARY FIGURE LEGENDS

**Supplementary Figure S1. Effect of cytokinin on STM levels. A,** Representative GFP images from *pTCSn::GFP* SAMs from plants grown under high light (HL; 170 μmol m^-2^ s^-1^) or low light (LL; 60 μmol m^-2^ s^-1^) conditions and of plants transferred from HL to darkness (D) for 48 h. Scale bar, 50 μm. **B,** GFP quantification from SAMs of *pTCSn::GFP* plants grown under HL or LL conditions. Plots show SAM measurements of plants grown as 3 independent batches normalized by the mean of the HL condition of each batch (HL: *n*=44, LL: *n*=41). Student’s *t*-test (*p*-values shown). **C,** GFP quantification from SAMs of *pTCSn::GFP* plants grown in HL and transferred to D or kept under HL for 48 h. Plots show SAM measurements of plants grown as 3 independent batches normalized by the mean of the HL condition of each batch (HL, *n*=44; 48 h D, *n*=45). Student’s *t*-test (*p*-values shown). **D,** Effect of cytokinin (BAP) application on the expression of the STM-VENUS reporter. SAMs of plants grown under HL conditions were excised and placed in medium supplemented or not with 500 nM BAP for the indicated times. Plots show SAM measurements of plants grown as 2 independent batches normalized by the mean of the uncut (0h) condition of each batch (*n*=14 for each of the indicated conditions). Different letters indicate statistically significant differences (Kruskal-Wallis with Dunn’s test; *p*<0.05). **E,** Effect of cytokinin (BAP) application on the activity of the *pTCSn::GFP* reporter. SAMs of plants grown under HL conditions were excised and placed in medium supplemented or not with 500 nM BAP for the indicated times. Plots show SAM measurements of plants grown as 2 independent batches normalized by the mean of the uncut (0h) condition of each batch (all 0 nM BAP conditions, *n*=12; 0h 500 nM BAP, *n*=12; 24 h 500 nM BAP, *n*=10; 48 h 500 nM BAP, *n*=10). Different letters indicate statistically significant differences (Kruskal-Wallis with Dunn’s test; *p*<0.05).

**Supplementary Figure S2. Effect of sugar on *STM* promoter activity. A,** Effect of light on the levels of soluble sugars in rosettes of *pSTM::STM-VENUS* plants grown in high light (HL) and transferred to darkness (D) or kept in HL for 48 h. Suc, sucrose; Tre6P, trehalose 6-phosphate; Glc, glucose; Fru, fructose. Plots show measurements of 6 whole rosettes from plants grown as one batch. Welch’s *t*-test (*p*-value shown). **B,** Effect of light on STM-VENUS levels in cut inflorescences. Inflorescences of *pSTM::STM-VENUS* plants grown under HL were cut and placed in medium without sugar for 48 h under HL (L) or dark (D) conditions, after which the SAMs were dissected and imaged (VENUS). Upper panel, representative STM-VENUS images of SAMs. Scale bar, 50 μm. Lower panel, plots showing SAM measurements of plants grown as one batch normalized by the mean of the uncut condition (uncut, *n*=7; 48 h L, *n*=7; 48 h D, *n*=7). Different letters indicate statistically significant differences (Kruskal-Wallis with Dunn’s test; *p*<0.05). **C,** Effect of sugar on STM-VENUS levels in cut inflorescences. Inflorescences of *pSTM::STM-VENUS* plants grown under HL condition were cut and placed under darkness for 48 h in medium with 2% sucrose or 1% sorbitol as osmotic control. SAMs were thereafter dissected and imaged (VENUS). Upper panel, representative STM-VENUS images of SAMs. Scale bar, 50 μm. Lower panel, plots showing SAM measurements of plants grown as one batch normalized by the mean of the uncut condition (uncut, *n*=7; 1% Sor, *n*=7; 2% Suc, *n*=7). Different letters indicate statistically significant differences (Kruskal-Wallis with Dunn’s test; *p*<0.05). The same batches of HL-grown uncut plants served as controls for the experiments shown in (**B)** and (**C)**. **D,** Effect of light on *STM* promoter activity in cut inflorescences. Inflorescences of *pSTM::STM-VENUS/pSTM::TFP-N7* plants grown under HL were treated as in (B) and dissected SAMs were imaged (TFP). Upper panel, representative *pSTM::TFP-N7* images of SAMs. Scale bar, 50 μm. Lower panel, plots showing SAM measurements of plants grown as 2 independent batches (except 48h L samples, which generated from a single batch) normalized by the mean of the *uncut* condition of each batch (uncut, *n*=17; 48 h L, *n*=7; 48 h D, *n*=14). Different letters indicate statistically significant differences (Kruskal-Wallis with Dunn’s test; *p*<0.05). **E,** Effect of sugar on *STM* promoter activity in cut inflorescences. Inflorescences of *pSTM::STM-VENUS/pSTM::TFP-N7* plants grown under HL were cut and placed under darkness for 48 h in medium with sucrose (Suc; 2% and 5%) or sorbitol (Sor; 1% and 2.5%) as osmotic control. SAMs were thereafter dissected and imaged (TFP). Upper panel, representative *pSTM::TFP-N7* images of SAMs. Scale bar, 50 μm. Lower panel, plots showing SAM measurements of plants grown as 2-3 independent batches (except 2.5% Sor and 5% Suc, which were grown as a single batch) normalized by the mean of the *uncut* condition of each batch (uncut, *n*=24; 1% Sor, *n*=14; 2% Suc, *n*=21; 2.5% Sor, *n*=7; 5% Suc, *n*=6). Different letters indicate statistically significant differences (Kruskal-Wallis with Dunn’s test; *p*<0.05).

**Supplementary Figure S3. Effect of ubiquitous SnRK1α1 overexpression on *STM* levels.** RT-qPCR analyses of *STM* in SAMs of control and *SnRK1α1-OE* plants grown under high light conditions (170 μmol m^-2^ s^-1^). Graphs correspond to the average of 3 independent samples, each consisting of a pool of 5 SAMs. Paired ratio *t*-test (*p*-value shown).

**Supplementary Figure S4. Effect of ubiquitous SnRK1α1 overexpression on the accumulation of soluble sugars**. Control and *SnRK1α1-OE* plants were grown under HL conditions and transferred to darkness (D) or kept under HL for 48 h. Plots show measurements of 6 whole rosettes from plants grown as one batch, with *SnRK1α1-OE* values expressed in comparison to the control. Suc, sucrose; Tre6P, trehalose 6-phosphate. Glc, glucose; Fru, fructose. Welch’s *t*-test (mutant *vs*. control for each condition; *p*-value shown).

**Supplementary Figure S5. Effect of SnRK1α depletion in the SAM on *STM* levels**. RT-qPCR analyses of STM in SAMs of control and *pSTM::amiRα* plants grown under high light conditions (170 μmol m^-2^ s^-1^). Two independent lines of *pSTM::amiRα-2 (amiRα-2#1* and *amiRα-2*#2) and *pSTM::amiRα-1 (amiRα-1#1* and *amiRα-1#2*) were used. Graph shows measurements from 2 independent samples, each consisting of a pool of 5 SAMs.

**Supplementary Figure S6. Effect of SnRK1α depletion in the SAM on organ boundary formation. A,** Representative meristems expressing *STM-VENUS* together with *pSTM::amiRα-1, pSTM::amiRα-2* or *pSTM::TFP-N7* as a control, and whose membranes were labelled with FM4-64. Left panels show the sum-slice projection of the FM4-64 signal and the right panels, two longitudinal sections showing the SAM-organ boundary of two stage-2 primordia. Note the reduced folding of the boundary and reduced bulging of the primordia in the *pSTM::amiRα-2* line. Scale bars, projections: 50 μm, sections: 20 μm. **B,** Folding angle of the boundary as a function of the size of the primordium in control plants (expressing *pSTM::TFP-N7*) as compared to *amiRα-2* (upper graph) or *amiRα-1* plants (lower graph). Plants in the *amiRα-2* experiment were grown as 3 independent batches. Control, *n*=126 primordia from 29 SAMs; *amiRα-2#1, n*=121 primordia from 25 SAMs; *amiRα-2#2, n*=120 primordia from 26 SAMs). Plants in the *amiRα-1* experiment were grown as 2 independent batches. Control, *n*=95 primordia from 22 SAMs; *amiRα-1#1, n*=84 primordia from 22 SAMs; *amiRα-1#2, n=87* primordia from 21 SAMs. Data were fitted using a linear regression (see Methods). #1 and #2 denote two independent lines for the indicated *amiRs*.

**Supplementary Figure S7. Silencing of *SnRK1α* in the SAM compromises meristem function and plant architecture. A,** Representative images of plants expressing *STM-VENUS* (control) together with *pSTM::amiRα-1* or *pSTM::amiRα-2* and grown under equinoctial conditions until the completion of flowering. **B-C**, Representative images of 22d-old plants of the most severely affected *amiRα* line (*amiRα-2*#1) showing activation of axillary meristems before flowering (red arrows in **B-C**) and termination of the main meristem (white arrow in **C**). **D,** Representative images of plants expressing *STM-VENUS* together with *pSTM::TFP-N7* (control), *pSTM::amiRα-1*, or *pSTM::amiRα-2*, and grown under short-day conditions for 3 weeks and then transferred to long-day conditions until the completion of flowering. #1 and #2 denote two independent lines for the indicated *amiRs*.

