## Supplementary Figures and Methods for "Sugar signaling modulates SHOOT MERISTEMLESS expression and meristem function in Arabidopsis"

### Supplementary Figure S1

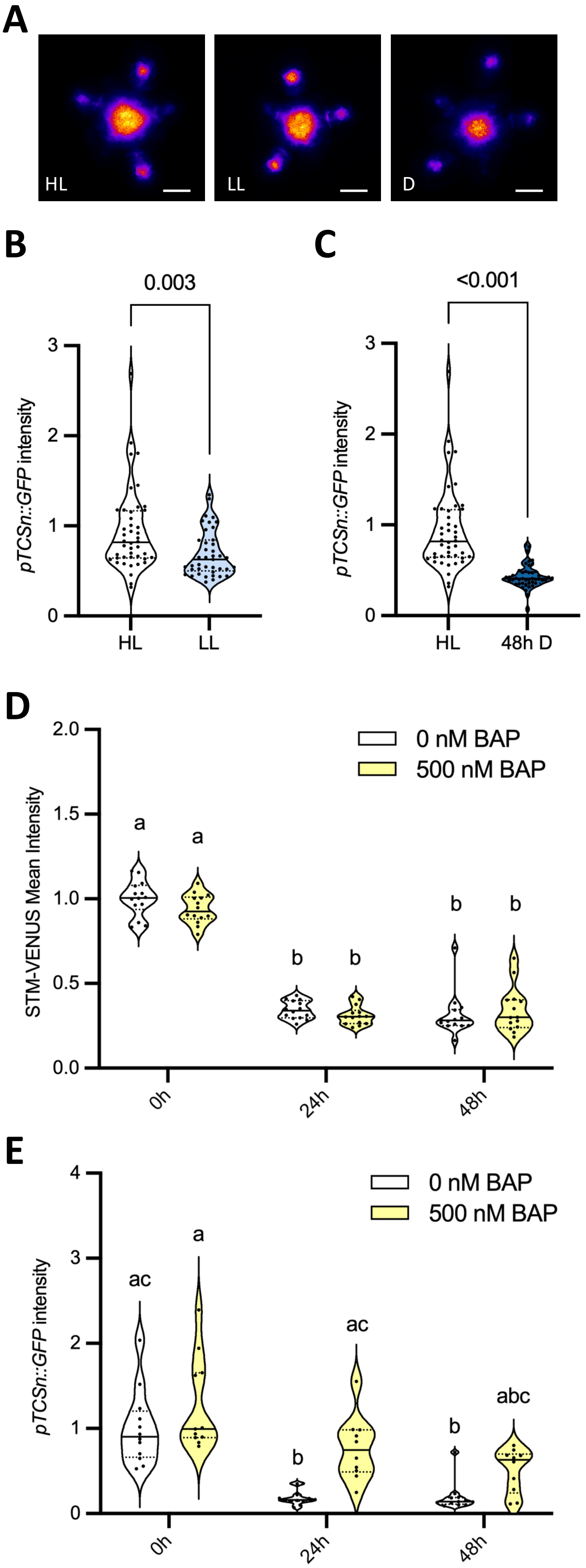

**Supplementary Figure S1. Effect of cytokinin on STM levels. A,** Representative GFP images from *pTCSn::GFP* SAMs from plants grown under high light (HL; 170 µmol $m^{-2}$ $s^{-1}$) or low light (LL; 60 µmol $m^{-2}$ $s^{-1}$) conditions and of plants transferred from HL to darkness (D) for 48 h. Scale bar, 50 µm. **B,** GFP quantification from SAMs of *pTCSn::GFP* plants grown under HL or LL conditions. Plots show SAM measurements of plants grown as 3 independent batches normalized by the mean of the HL condition of each batch (HL: $n$=44, LL: $n$=41). Student's $t$-test ($p$-values shown). **C,** GFP quantification from SAMs of *pTCSn::GFP* plants grown in HL and transferred to D or kept under HL for 48 h. Plots show SAM measurements of plants grown as 3 independent batches normalized by the mean of the HL condition of each batch (HL, $n$=44; 48 h D, $n$=45). Student's $t$-test ($p$-values shown). **D,** Effect of cytokinin (BAP) application on the activity of the *STM-VENUS* reporter. SAMs of plants grown under HL conditions were excised and placed in medium supplemented or not with 500 nM BAP for the indicated times. Plots show SAM measurements of plants grown as 2 independent batches normalized by the mean of the uncut (0h) condition of each batch ($n$=14 for each of the indicated conditions). Different letters indicate statistically significant differences (Kruskal-Wallis with Dunn´s test; $p$<0.05). **E,** Effect of cytokinin (BAP) application on the activity of the *pTCSn::GFP* reporter. SAMs of plants grown under HL conditions were excised and placed in medium supplemented or not with 500 nM BAP for the indicated times. Plots show SAM measurements of plants grown as 2 independent batches normalized by the mean of the uncut (0h) condition of each batch (all 0 nM BAP conditions, $n$=12; 0h 500 nM BAP, $n$=12; 24 h 500 nM BAP, $n$=10; 48 h 500 nM BAP, $n$=10). Different letters indicate statistically significant differences (Kruskal-Wallis with Dunn´s test; $p$<0.05).

### Supplementary Figure S2

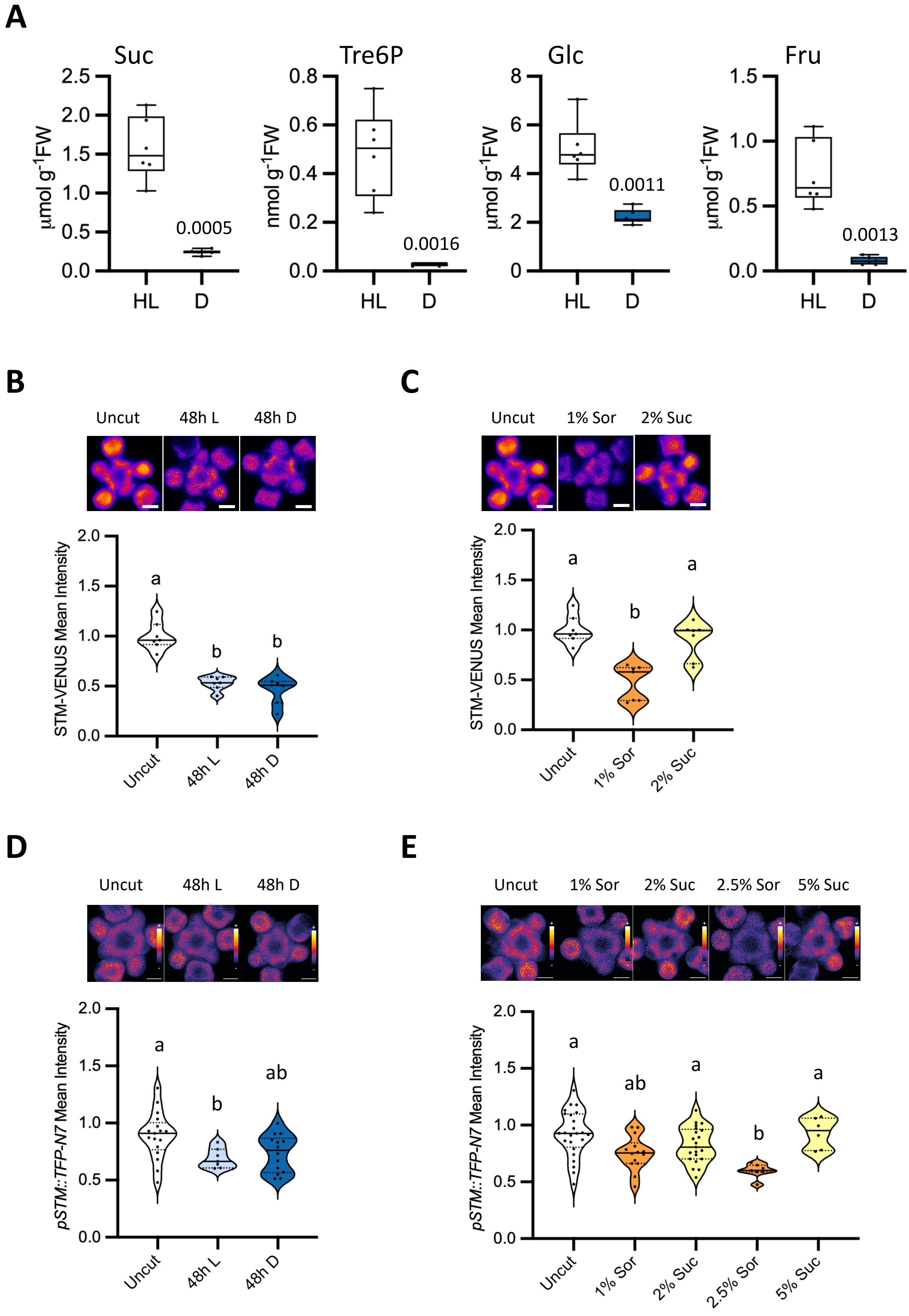

**Supplementary Figure S2. Effect of sugar on *STM* promoter activity. A,** Effect of light on the levels of soluble sugars in rosettes of *pSTM::STM-VENUS* plants grown in high light (HL) and transferred to darkness (D) or kept in HL for 48 h. Suc, sucrose; Tre6P, trehalose 6-phosphate; Glc, glucose; Fru, fructose. Plots show measurements of 6 whole rosettes from plants grown as one batch. Welch´s *t*-test (*p*-value shown). **B,** Effect of light on STM-VENUS levels in cut inflorescences. Inflorescences of *pSTM::STM-VENUS* plants grown under HL were cut and placed in medium without sugar for 48 h under HL (L) or dark (D) conditions, after which the SAMs were dissected and imaged (VENUS). Upper panel, representative STM-VENUS images of SAMs. Scale bar, 50 µm. Lower panel, plots showing SAM measurements of plants grown as one batch normalized by the mean of the uncut condition (uncut, *n*=7; 48 h L, *n*=7; 48 h D, *n*=7). Different letters indicate statistically significant differences (Kruskal-Wallis with Dunn´s test; $p<0.05$). **C**, Effect of sugar on STM-VENUS levels in cut inflorescences. Inflorescences of *pSTM::STM-VENUS* plants grown under HL condition were cut and placed under darkness for 48 h in medium with 1% sucrose or 2% sorbitol as osmotic control. SAMs were thereafter dissected and imaged (VENUS). Upper panel, representative STM-VENUS images of SAMs. Scale bar, 50 µm. Lower panel, plots showing SAM measurements of plants grown as one batch normalized by the mean of the uncut condition (uncut, *n*=7; 1% Sor, *n*=7; 2% Suc, *n*=7). Different letters indicate statistically significant differences (Kruskal-Wallis with Dunn´s test; $p<0.05$). The same batches of HL-grown uncut plants served as controls for the experiments shown in (B) and (C). **D**, Effect of light on STM promoter activity in cut inflorescences. Inflorescences of *pSTM::STM-VENUS/pSTM::TFP-N7* plants grown under HL were treated as in (B) and dissected SAMs were imaged (TFP). Upper panel, representative pSTM::TFP-N7 images of SAMs. Scale bar, 50 µm. Lower panel, plots showing SAM measurements of plants grown as 2 independent batches (except 48h L samples, which generated from a single batch) normalized by the mean of the uncut condition of each batch (uncut, n=17; 48 h L, n=7; 48 h D, n=14). Different letters indicate statistically significant differences (Kruskal-Wallis with Dunn´s test; $p<0.05$). **E**, Effect of sugar on *STM* promoter activity in cut inflorescences. Inflorescences of *pSTM::STM-VENUS;pSTM::TFP-N7* plants grown under HL were cut and placed under darkness for 48 h in medium with sucrose (Suc; 2% and 5%) or sorbitol (Sor; 1% and 2.5%) as osmotic control. SAMs were thereafter dissected and imaged (TFP). Upper panel, representative *pSTM::TFP-N7* images of SAMs. Scale bar, 50 µm. Lower panel, plots showing SAM measurements of plants grown as 2-3 independent batches (except 2.5% Sor and 5% Suc, which were grown as a single batch) normalized by the mean of the *uncut* condition of each batch (uncut, *n*=24; 1% Sor, *n*=14; 2% Suc, *n*=21; 2.5% Sor, *n*=7; 5% Suc, *n*=6). Different letters indicate statistically significant differences (Kruskal-Wallis with Dunn´s test; $p<0.05$).

### Supplementary Figure S3

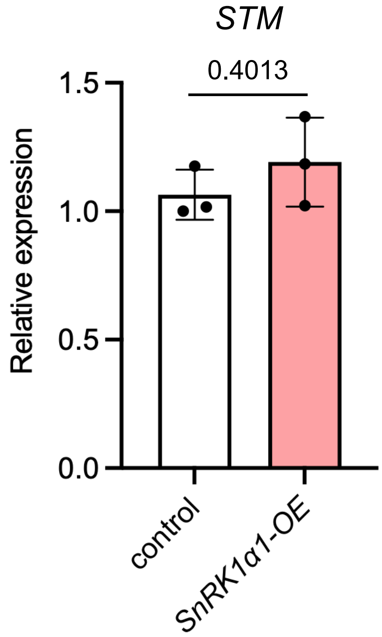


**Supplementary Figure S3. Effect of ubiquitous SnRK1α1 overexpression on *STM* levels.** RT-qPCR analyses of *STM* in SAMs of control and *SnRK1α1-OE* plants grown under high light conditions (170 µmol $m^{-2}$ $s^{-1}$). Graphs correspond to the average of 3 independent samples, each consisting of a pool of 5 SAMs. Paired ratio *t*-test (*p*-value shown).

#### Supplementary Figure S4

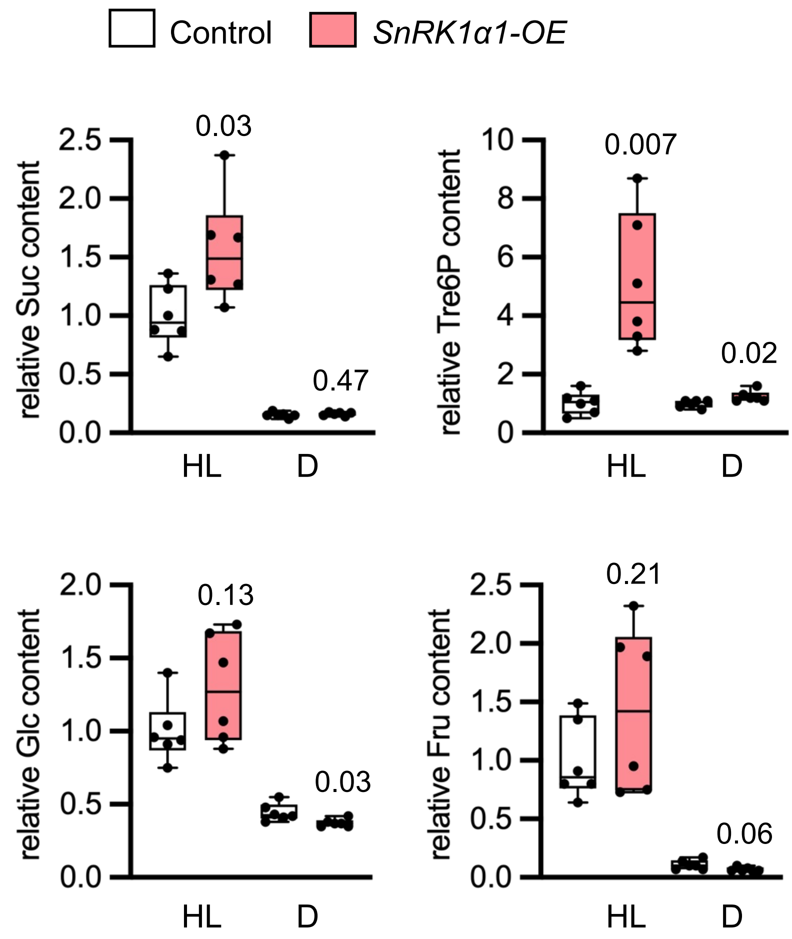


**Supplementary Figure S4. Effect of ubiquitous SnRK1α1 overexpression on the accumulation of soluble sugars**. Control and *SnRK1α1-OE* plants were grown under HL conditions and transferred to darkness (D) or kept under HL for 48 h. Plots show measurements of 6 whole rosettes from plants grown as one batch, with *SnRK1α1-OE* values expressed in comparison to the control. Suc, sucrose; Tre6P, trehalose 6-phosphate. Glc, glucose; Fru, fructose. Welch´s *t*-test (mutant *vs*. control for each condition; *p*-value shown).

#### Supplementary Figure S5

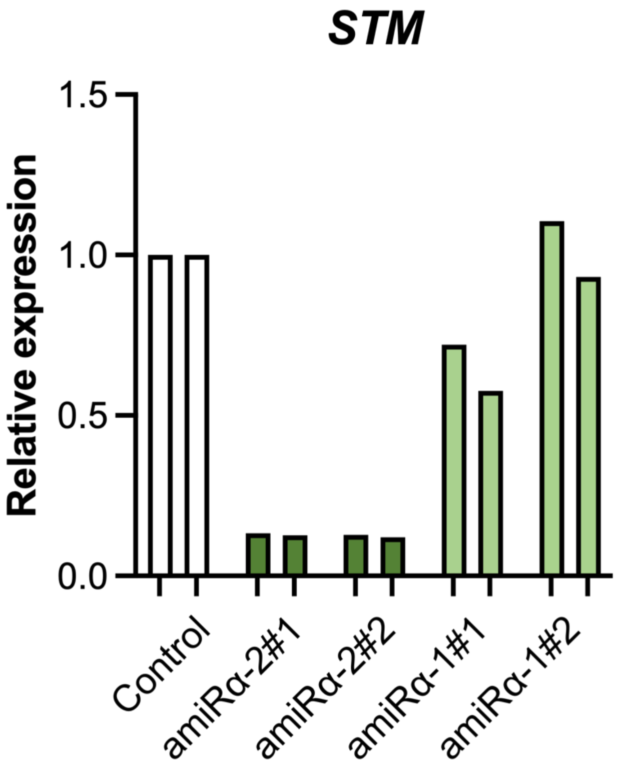


**Supplementary Figure S5. Effect of SnRK1α depletion in the SAM on *STM* levels**. RT-qPCR analyses of *STM* in SAMs of control and *pSTM::amiRα* plants grown under high light conditions (170 µmol $m^{-2}$ $s^{-1}$). Two independent lines of *pSTM::amiRα-2* (*amiRα-2*#1 and *amiRα-2*#2) and *pSTM::amiRα-1* (*amiRα-1*#1 and *amiRα-1*#2) were used. Graph shows measurements from 2 independent samples, each consisting of a pool of 5 SAMs.

### Supplementary Figure S6

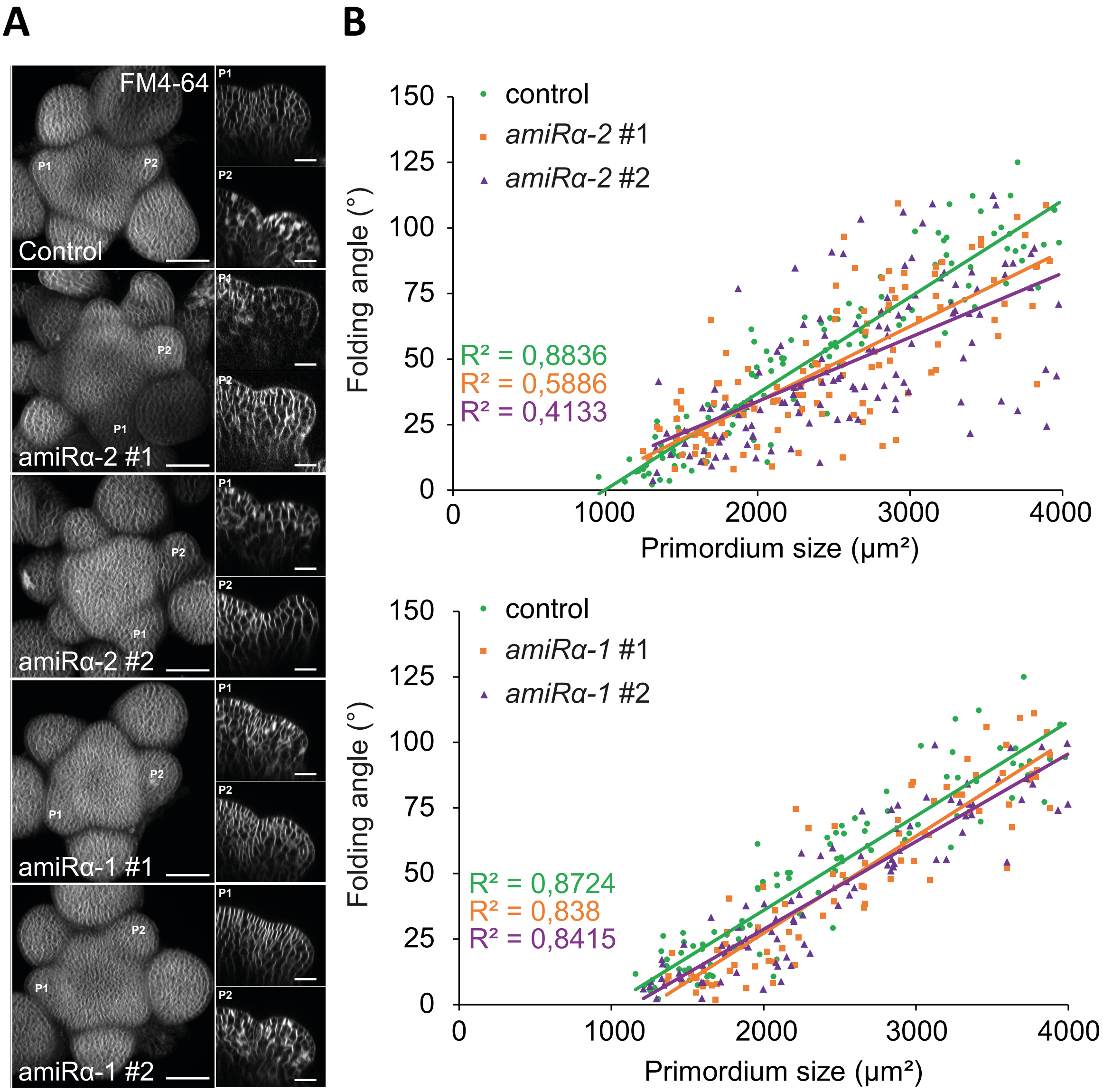


**Supplementary Figure S6. Effect of SnRK1α depletion in the SAM on organ boundary formation. A,** Representative meristems expressing *STM-VENUS* together with *pSTM::amiRα-1, pSTM::amiRα-2* or *pSTM::TFP-N7* as a control, and whose membranes were labelled with FM4-64. Left panels show the sum-slice projection of the FM4-64 signal and the right panels, two longitudinal sections showing the SAM-organ boundary of two stage-2 primordia. Note the reduced folding of the boundary and reduced bulging of the primordia in the *pSTM::amiRα-2* line. Scale bars, projections: 50 µm, sections: 20 µm. **B,** Folding angle of the boundary as a function of the size of the primordium in control plants (expressing *pSTM::TFP-N7*) as compared to *amiRα-2* (upper graph) or *amiRα-1* plants (lower graph). Plants in the *amiRα-2* experiment were grown as 3 independent batches. Control, *n*=126 primordia from 29 SAMs; *amiRα-2*#1, *n*=121 primordia from 25 SAMs; *amiRα-2*#2, *n*=120 primordia from 26 SAMs). Plants in the *amiRα-1* experiment were grown as 2 independent batches. Control, *n*=95 primordia from 22 SAMs; *amiRα-1*#1, *n*=84 primordia from 22 SAMs; *amiRα-1*#2, *n*=87 primordia from 21 SAMs. Data were fitted using a linear regression (see Methods).

### Supplementary Figure S7

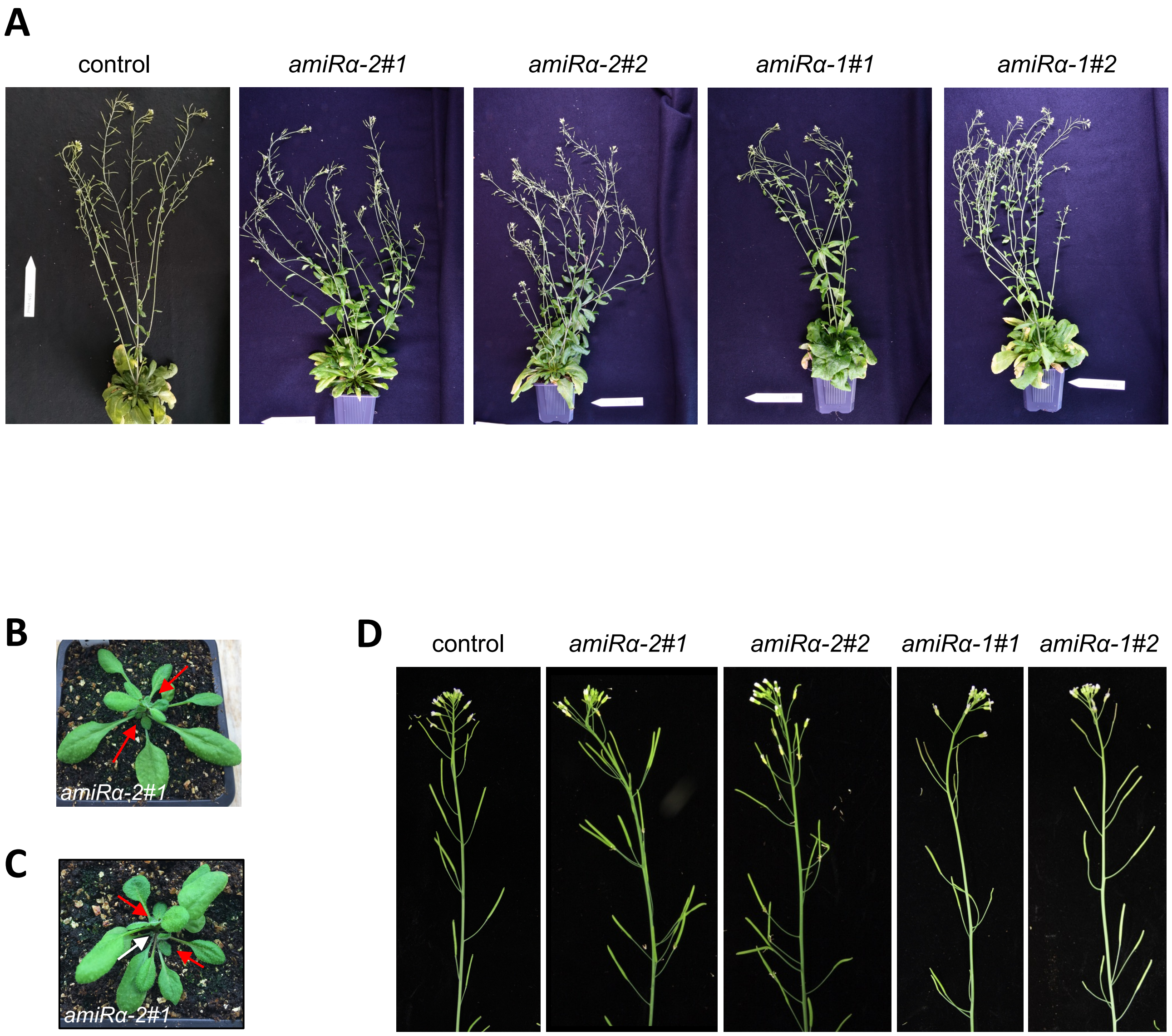


**Supplementary Figure S7. Silencing of *SnRK1α* in the SAM compromises meristem function and plant architecture. A,** Representative images of control plants and two independent *amiRα-2* and *amiRα-1* lines grown under equinoctial conditions until the completion of flowering. **B-C**, Representative images of 22d-old plants of the most severely affected *amiRα* line (*amiRα-2*#1) showing activation of axillary meristems before flowering (red arrows in **B-C**) and termination of the main meristem (white arrow in **C**). **D,** Representative images of control plants (expressing *pSTM::TFP-N7*) and two independent *amiRα-2* and *amiRα-1* lines grown under short-day conditions for 3 weeks and then transferred to long-day conditions until the completion of flowering.

**SUPPLEMENTARY INFORMATION**

**Plant material**

All *Arabidopsis thaliana* (L.) Heynh. plants used here are in the Columbia (Col-0) background. The *pSTM::STM-VENUS* line (*STM-VENUS*) was generated by transforming Col-0 plants with the plasmid described by Heisler and colleagues (1, 2). The *pTCSn::GFP* line was provided by Bruno Müller (3). The *SnRK1α1-GFP* [*pSnRK1α1::SnRK1α1-GFP::terSnRK1α1/snrk1α1-3*; (4)] and *SnRK1α1-OE* [*35S::SnRK1α1*; (5)] lines were previously described. For expression of STM-VENUS in the *SnRK1α1-OE* background, the *SnRK1α1-OE* and *STM-VENUS* lines were crossed, and homozygous progeny was selected on kanamycin and BASTA. For generating the *pSTM::STM-VENUS/pSTM::TFP-N7* line, the 5.7 kb promoter of *STM* was amplified by PCR and introduced in a pDONR-P4-P1R plasmid by Gateway recombination cloning (6). A triple LR reaction was then performed between this vector, a TFP-N7 recombined with a pDONR221, a Nos-T7 terminator recombined with a pDONR-P2R-P3, and a pH7m34GW destination vector. The resulting gene construct was then introduced into *Arabidopsis* plants already expressing the *STM-VENUS* reporter using *Agrobacterium*. Transformants were selected based on the hygromycin resistance of the pH7m34GW vector. For generating the *pSTM::amiRα* lines, two amiRNAs targeting both *SnRK1α1* and *SnRK1α2* (*amiRα-1* and *amiRα-2*) were designed using the WMD3 Web microRNA designer tool (7) and introduced into the primary *miRNA319a* (At4g23713) backbone in the pHBT95 vector (8). The resulting primary *miRNA319a*, harboring the *amiRα-1* or *amiRα-2* sequence, was used to generate the corresponding *amiRα-1* and *amiRα-2* pENTRY 1_2 constructs and recombined with *pSTM* (5.7 kb) pENTRY 4_1r and *NOS-T7*-pENTRY 2r_3 into pK7m34GW (9). The resulting construct was transformed into the *STM-VENUS* line and single-insertion homozygous progeny was selected on kanamycin.

**Plant growth conditions**

Seeds were sown in excess into individual pots (7 x 7 x 6 cm, W/L/H) with soil (Levington F2) that were thereafter kept in a 4°C room for three days before being transferred to growth cabinets. The pots were covered with transparent plastic lids during the first five days and the plants were thinned out seven days after sowing to leave one plant per pot. For most experiments, seedlings were initially grown in short-day conditions (8 h/16 h light/dark period) for 3-4 weeks and then transferred to continuous light (24 h light, temperature: 22°C) and kept under an irradiance of 60 (LL) or 170 (HL) µmol $m^{-2}$ $s^{-1}$, provided by white fluorescent lamps. Unless otherwise indicated, plants were grown in HL conditions. For dark treatment, plants grown under HL were put into the dark for 24, 48 or 72 h. For the assays with excised inflorescences and STM-VENUS imaging of *amiRα* lines, seedlings were grown in short-day conditions (8 h/16 h light/dark period) for 3-4 weeks and then transferred to long-day conditions (16 h/8 h light/dark period; 22°C/18°C).

**Protein extraction, quantification, and immunoblotting**

For extraction of total protein for the immunoblot analyses, one leaf or complete inflorescence meristems (five per replicate) were ground in liquid $N_2$. Finely ground tissue (approximately 30-40 mg per sample) was extracted in 1.5 volumes of buffer [150 mM NaCl, 1% (v/v) Triton-X, 50 mM Tris-HCl (pH 8.0), 3 mM DTT, supplemented with 50 μM MG132, 1:20 Complete EDTA-free Protease Inhibitor Cocktail (Roche), and 1:500 Phosphatase Inhibitor Cocktails 2 and 3 (Sigma-Aldrich; www.sigmaaldrich.com)]. Homogenates were cleared by centrifugation at 21130 x g for 15 minutes at 4°C, and supernatants were used for total protein quantification (Pierce 660 nm protein assay reagent; www.thermofisher.com). Single use aliquots of 40 μg total protein were prepared and denatured in Laemmli buffer(12) before storage at -20°C. Total protein samples were separated by SDS-PAGE in 10% acrylamide gels, transferred to PVDF membranes (wet transfer at 4°C, 100 V, 70 min) and probed using 1:1000 dilutions of STM, p-AMPK, SnRK1α1 or TUBULIN antibodies (see Table S1 for further details).

Chemiluminescent detection was performed using a 1:10,000 dilution of Peroxidase AffiniPure goat anti-rabbit IgG (H+L) (Jackson ImmunoResearch; www.jacksoniummuno.com), and SuperSignal West Femto Maximum Sensitivity Substrate (Thermo Scientific).

Band intensity was quantified using ImageJ. The bands of interest were selected with the rectangle tool and the corresponding intensity peaks were plotted. The peaks were separated using the line tool after which their area was measured. For quantifying STM-VENUS and SnRK1α1 amounts, band intensity values were normalized to the values corresponding to their respective Rubisco band in the Ponceau-stained membrane. For quantifying SnRK1α phosphorylation, band intensity values of the phospho-SnRK1α immunoblot were normalized to the values corresponding to their respective SnRK1α1 immunoblot.

**RNA extraction, cDNA synthesis and qRT-PCR**

Total RNA was extracted from approximately 30-40 mg of finely ground tissue, with the Plant Nucleospin RNA extraction kit (Macherey-Nagel; www.mn-net.com), according to the manufacturer's instructions. DNAse-treated RNA (1 μg) was used for synthesis of cDNA libraries, using SuperScript III Reverse Transcriptase (Life Technologies;www.thermofisher.com).

qRT-PCR analyses were performed in 384-well reaction plates using the QuantStudio™ 7 Flex Real- Time PCR System (Thermo Fisher). The reactions were prepared in a total volume of 10 μL containing 1 μL of cDNA (diluted 1:10) corresponding to 2.5 ng of RNA, 5 μL of iTaq

Universal SYBR Green Supermix (BioRad www.bio-rad.com) and 0.8 µL of each gene-specific 5 µM primer. No-template and -RT controls were included for each gene in comparative gene expression analyses.

The $2^{-\Delta\Delta Ct}$ method was used for relative quantification. Expression values were normalised to the geometric mean of Ct values obtained for the following reference genes: *UBQ10* (At4g05320) and *UBC21* (At5g25760). Expression of reference genes, *STM* (At1g62360), *HB25* (At5g65410), *AIL7* (At5g65510), *DIN10* (At5g20250), *DRM2* (At2g33830) and *SEN5* (At3g15450) was assessed using the primers described in Table S1.

**Sugar measurements**

Soluble sugars were extracted from aliquots (15-20 mg) of frozen tissue powder using chloroform-methanol as described in (13). Sucrose, glucose and fructose were quantified by high-performance hydrophilic interaction chromatography on a 150 x 2.1 mm Luna Omega Sugar (Phenomenex Inc.; Aschaffenburg, Germany; www.phenomenex.com) column, fitted with a 4 x 2.0 mm Security Guard Cartridge (Phenomenex) and coupled to a QTrap 5500 triple quadrupole mass spectrometer (AB Sciex, Foster City, USA; sciex.com). Samples were spiked with $^{13}C$-labelled internal standards for correction of ion suppression and other matrix effects (14). The column was maintained at 25°C and equilibrated with 90 % (v/v) acetonitrile, 5 % (v/v) isopropanol, 5 % (v/v) water (eluent A) before injection of samples (1 µl). Sugars were eluted (flow rate 0.25 ml min-1) with the following gradient, obtained by mixing eluent A with eluent B [5 % (v/v) acetonitrile, 25 % (v/v) isopropanol, 70 % (v/v) water): 0 – 2.5 min [10 % B]; 2.5 - 25 min [10 - 30 % B]; 25 – 30.5 min [30 % B]; 30.5 – 31 min [30 – 10 % B]; 31 – 40 min [10 % B]. Sugars were quantified by tandem mass spectrometry using the parent/product ion transitions as described in (14). Tre6P was quantified in chloroform-methanol extracts as described in (13) with modifications as described in (15).

**Y2H assays**

For Y2H assays, a pGBKT7 construct harboring the *SnRK1α1* coding sequence had previously been generated (16). The *STM* coding sequence was amplified from seedling cDNA and cloned into pDONR221 and the resulting pENTRY-STM construct was recombined into a Gateway-compatible pDEST-GADT7 vector (17). Y2H assays were performed as described (18). To test for interaction between SnRK1α1 and STM, yeast cells were co-transformed with the pGBKT7-SnRK1α1 construct, harboring the GAL4 DNA-binding domain (BD), and the pDEST-GADT7-STM construct, harboring the GAL4 activation domain (AD). As controls, pGBKT7-SnRK1α1 subunit and pDEST-GADT7-STM constructs were co-transformed with the respective complementary empty vectors. Yeast growth was assessed in low-stringency media [-Leucine (L)/-Tryptophan(W)] for selection of co-transformants, and in higher-stringency media [-L/-W/-Histidine (H)/] for selection of interactors.

**Protoplast assays**

Protoplast transient expression assays were carried out as previously described, using freshly isolated mesophyll cells from mature fully expanded Arabidopsis leaves (19, 20). For the expression of SnRK1α1 with a C-terminal HA tag, a previously described pHBT95-based construct was used (19). For expressing STM with a C-terminal GFP tag, a pENTRY-STM construct was recombined with the p2GWC7 vector (9). pHBT95-SnRK1α1-HA and p2GWC7-STM-GFP were used in a 1:1 ratio to co-transfect 1 ml of mesophyll cell protoplasts. Following transfection, protoplasts were incubated for 16h under light conditions (25-30 μmol $m^{-2}$ $s^{-1}$ and 23°C), and then harvested by centrifugation and flash frozen.

Frozen cell pellets were lysed in 500 µl of lysis buffer [150 mM NaCl, 0.5% (v/v) Triton-X, 50 mM Tris-HCl (pH 8.0), 3 mM DTT, supplemented with 50 µM MG132, 1:20 Complete EDTA-free Protease Inhibitor Cocktail (Roche), and 1:500 Phosphatase Inhibitor Cocktails 2 and 3 (Sigma-Aldrich)]. The cleared lysate was incubated with 30 µl of super-paramagnetic µMAC beads coupled to a monoclonal anti-GFP antibody (Miltenyi Biotec) for 2h at 4°C with gentle rocking. Purified immunocomplexes were eluted in Laemmli buffer, boiled, run in an 10% SDS-PAGE gel, transferred to PVDF membranes (wet transfer at 4°C, 100 V, 70 min) and analysed by immunoblotting using antibodies against GFP (1/1000, Roche, #11814460001), HA (1/1000, Roche, #11867423001), and SnRK1α1 (21), used at a 1:2000 dilution.

For immunodetection, secondary antibodies conjugated with horseradish peroxidase (AffiniPure goat anti-rabbit (#111035144) or anti-rat IgG (H+L) (#112035167); Jackson ImmunoResearch; www.jacksoniummuno.com) were used at 1:10000 dilutions. Chemiluminescence-based detection of peroxidase activity was performed using a SuperSignal West Femto Maximum Sensitivity Substrate (Thermo Scientific).
